# Generation of a Three-dimensional Human Neurovascular Unit Model in a Microfluidic Chip

**DOI:** 10.1101/2025.04.13.648605

**Authors:** Boning Qiu, Sara Pompe, Katerina T. Xenaki, Paul M.P. van Bergen en Henegouwen, Sabrina Oliveira, Enrico Mastrobattista, Massimiliano Caiazzo

## Abstract

**Background:** The well-functioning of the neurovascular unit (NVU) is supported by the 3D brain physiological microenvironment that allows for extensive neural-neural and neural-vascular interactions. This microenvironment is normally hard to create in traditional *in-vitro* models such as the transwell model. Organ-on-a-chip (OOC) emerges as advanced model systems by providing better physiological microenvironments. However, NVU modeling in many chip platforms has not met a full 3D condition for neural cultures.

**Methods:** Here, we describe a novel NVU model generated in a microfluidic chip that reproduces the neural-neural and neural-vascular interactions in a full-3D format. The model features an extracellular matrix (ECM) environment that supports both a perfused brain endothelial vessel and 3D cultured neural cells (astrocytes and neurons) beside the tube. Culture conditions were comprehensively optimized for better endothelial tube integrity as well as ECM gel longevity. The model was used to model neuroinflammation-induced brain tube disruption and immune cell extravasation. Furthermore, as a drug testing platform, the model was explored for brain endothelial transcytosis of the heparin-binding EGF-like growth factor (HB-EGF) targeted nanobodies and the data was compared to a parallel transwell model.

**Results:** Immunofluorescent staining confirmed the expression of endothelial junctional proteins, as well as astrocytic and neuronal markers. The perfused brain endothelial tube exhibited resistance to paracellular leakage of 20 kDa FITC-dextran. Astrocytes and neurons growing in ECM gel developed extensive neural network and showed spontaneous neuronal firing. The neural-vascular interactions were formed through astrocyte migration and axonal outgrowth in the ECM gel towards the tube. Exposure to neuroinflammatory cytokines disrupted the tube barrier, resulting in increased barrier leakage and the recruitment of peripheral blood mononuclear cells (PBMCs) as well as their extravasation. Owing to full-3D model design, endothelial transcytosis and abluminal distribution of the fluorescently labeled HB-EGF targeting Nbs can be clearly visualized in situ. Compared to a transwell model counterpart, the NVU chip model performed better in revealing the binding and transcytosis specificity of the targeted nanobodies.

**Conclusions:** We demonstrate improved physiological relevance in this full-3D NVU-on-a-chip model. The model could become a faithful platform for NVU research under both healthy and diseased conditions, and can be used as a reliable drug testing platform that aims at developing novel brain-targeted therapeutics.

## Introduction

The blood–brain barrier (BBB), consisting of brain endothelial cells, basement membrane, pericytes and astrocyte end-feet, is a highly integrated physiological structure that exquisitely mains brain homeostasis [1–3]. On the other hand, the BBB is also functionally coupled with surrounding neurons, all of which are together termed as neurovascular unit (NVU), a basic functioning unit of the brain [1, 3, 4]. The advantages of having reduced model complexities and easy experimental setups make *in-vitro* models suitable for unraveling multi-faceted roles of the NVU in contributing to brain homeostasis or dysregulations and for prescreening drugs that specifically target and bypass the BBB to treat pathological neural cells [5]. While animal cells bare substantial species gaps in neurophysiology [6–12], accessibility to human cells grants the *in-vitro* NVU models better authenticity in modeling human brain physiology and in predicting for clinical translation of the therapeutical molecules being tested [13]. Nonetheless, most of the *in-vitro* NVU models today still face difficulties in reproducing the complexity of the brain microenvironment, which makes them sub-qualified substitutes to the animal models in some cases.

Recently, microfluidic technology has greatly revolutionized *in-vitro* modeling by introducing the concept of organ-on-a-chip (OOC) [14, 15]. Microfluidic chips enable the modeling of a more physiological microenvironment of the NVU, including the formation of perfused brain endothelial vessels and the incorporation of related neural cell types in 3D extracellular matrix (ECM) gel [14–17]. The presence of these *in-vivo* like conditions was found to induce a more physiologically relevant phenotype for both the brain endothelial cells and the neural cells [18–41]. Many of the chip designs for modeling various cell barriers incorporate parallelled channels that are convenient for spatially segregating the vasculatures and the perivascular cells while still allowing for their intercommunications [40, 42, 43]. Advanced chip designs further permit a membrane-free environment for growing the vasculatures, which greatly promotes the vascular-ECM and vascular-stromal cell interactions [30, 44, 45].

So far, BBB or NVU models developed on those multi-channel chips, despite the generation of perfused brain endothelial vessels, did not provide a proper 3D ECM environment for the neural culture [33, 34, 40, 46–48]. These models failed to achieve the well-developed 3D neural network as well as proper neural-vascular interactions, which are important anatomical foundations of the functioning of the NVU [1]. Strikingly, some models even subjected the adherent neural culture directly under medium perfusion [33, 34, 40], which did not faithfully recapitulate physiological relevance. In this study, we reported an optimized NVU modeling protocol in a 3-lane chip platform for further improving the model’s physiological resemblance. Our full-3D NVU model consists of a perfused brain microvascular endothelial tube in the top channel and neural cells (astrocytes and neurons) embedded in Matrigel in the bottom channel, where the two sections are separated by a middle channel collagen gel. The double-gel structure allows for a stable endothelial tube culture and meanwhile permits the 3D neural network formation, followed by vascular-neural contact through the astrocyte migration and the axonal outgrowth. We showed the model’s value for mimicking neuroinflammation-induced BBB disruption and immune cell extravasation. Besides, we used this model for testing BBB shuttle nanobodies (Nbs). The ability of two previously identified Heparin-binding EGF-like growth factor (HB-EGF) targeted Nbs to transcytose the BBB were revealed in this model. Notably, when using fluorescently labeled Nbs, the 3D neural environment allowed for easy in-situ visualization of the Nb transcytosis and distribution. With better physiological relevance, our full-3D NVU model could become a faithful *in-vitro* platform to study the basic functions of the human brain, and to be used as a reliable drug prescreening platform that aims at developing the brain-targeting therapeutics.

## Methods

### Cell culture

Primary Human Brain Microvascular Endothelial Cells (hBMECs) (ACBRI 376) were purchased from Cell Systems. Cells were maintained in culture flask coated with 50 µg/ml collagen-IV (C5533, Sigma-Aldrich) and 25 µg/ml fibronectin (F4759, Sigma-Aldrich) in MV-2 medium (C-2212, Promocell) supplemented with 1× Antibiotic Antimycotic reagent (A5955, Sigma-Aldrich) and cultured in 5% CO_2_ humidified incubator at 37 ℃. Medium was refreshed every two days. Upon reaching confluency around 70%, cells were detached with 0.025% trypsin EDTA (#25300054, Thermo Fisher) and sub-cultured at ratio of 1:3-1:6 in new coated flasks. HBMECs were used between passage 4 and 9.

Primary human astrocytes (#1800) were purchased from ScienCell. Cells were maintained in culture flask coated with hESC-qualified Matrigel (#354277, Corning) in 5% CO_2_ humidified incubator at 37 ℃. Astrocyte growth medium (AGM, #1801, ScienCell) supplemented with 1× Antibiotic Antimycotic was used. Medium was refreshed every two days. Upon reaching confluency around 90%, cells were detached with 0.025% trypsin EDTA and sub-cultured at the density of 5,000 cells/cm^2^ in new coated flasks. Astrocytes were used between passage 4 and 9.

LUHMES cells (CRL-2927), a human mesencephalic neural cell line, were purchased from ATCC. Cells were maintained in culture flask coated with 50 µg/mL poly-L-ornithine (PLO, P4957, Sigma-Aldrich) and 2 µg/ml fibronectin (F4759, Sigma-Aldrich) and cultured in 5% CO_2_ humidified incubator at 37℃. LUHMES maintenance medium consists of advanced DMEM/F12 (#12634010, Thermo Fisher), 0.5× GlutaMAX (#35050061, Thermo Fisher), 1× N2 supplement (#17502048, Thermo Fisher), 10 ng/ml FGF-2 (#100-18b, Peprotech) and 1× Antibiotic Antimycotic. For sub-culture, upon reaching 70-80% confluency, cells were detached with 0.05% trypsin EDTA (#25300054, Thermo Fisher) and split at ratio of 1:10 in new coated flasks. Medium was renewed every other day. LUHMES cells were used between passage 5 and 12. To differentiate LUHMES cells into DA neurons, a modified two-step protocol was derived based on the original protocol [49]. Briefly, LUHMES sub-culture at 40-50% confluency weas subjected to pre-differentiation medium which consisted of advanced DMEM/F12, 0.5× GlutaMAX, 1× N2 supplement, 1× Antibiotic Antimycotic, 1mM db-cAMP (D0627, Sigma-Aldrich), 20 ng/ml BDNF (#450-02, Peprotech), 20 ng/ml GDNF (#450-10, Peprotech), 20 ng/ml TGF-β3, 0.2 µM ascorbic acid (A4403, Sigma-Aldrich) and 2 µg/ml Doxycycline (D3072, Sigma-Aldrich). Medium was not changed for two days. For final differentiation, the cells were detached, counted, pelleted, and resuspend in final differentiation medium for re-plating either in 3D gel or in well plate. The final differentiation medium consists of NeuroBasal (#21103049, Thermo Fisher), 1× GlutaMAX, 1× B27 supplement (#12587010, Thermo Fisher), 1× Antibiotic Antimycotic, 1mM db-cAMP, 20 ng/ml BDNF, 20 ng/ml GDNF, 0.2 µM ascorbic acid, 2 µg/ml Doxycycline and 1 µM DAPT (D5942, Sigma-Aldrich).

Peripheral blood mononuclear cells (PBMCs) were isolated from healthy donors at the University Medical Center Utrecht (UMCU). PBMCs were maintained as suspension in culture flask in RPMI 1640 medium (R8758, Sigma-Aldrich) supplemented with 2.5% (v/v) human serum (H6914, Sigma-Aldrich), 0.1% (v/v) 2-mercaptoethanol (#21985023, Thermo Fisher), 50 IU/ml IL-2 (#200-02, Peprotech), 1× Antibiotic Antimycotic and cultured in 5% CO_2_ humidified incubator at 37 ℃. Freshly isolated PBMCs were used within 5 days.

### OrganoPlate NVU model

The 3-lane OrganoPlate (#4004-400B, MIMETAS) with 400 µm × 220 µm (w × h) channels and 100 µm × 55 µm (w × h) Phaseguide^TM^ was used for NVU modeling. The tri-seeding method was adapted from the original protocol provided by the MIMETAS company. Briefly, neutralized rat collagen-I (#354249, Corning) of 5 mg/ml was prepared on ice by mixing with 1 M HEPES and 37 g/l NaHCO_3_ at 8:1:1 (v/v/v) ratio followed by subsequent PBS dilution. 3 µl gel was dispensed into the middle channel of the chips and the plate was placed in the in 5% CO2 humidified cell incubator at 37℃ for 20 min. ES-qualified Matrigel (#354277, Corning) was diluted on ice with LUHMES pre-differentiation medium (see LUHMES culture section) to reach 5 mg/ml. Single astrocytes and 2- day pre-differentiated LUHMES cells were prepared and resuspend with Matrigel to achieve desired density (LUHMES: 20,000 cells/µl gel; astrocyte: 5,000 cells/µl gel). 3µl of the ice-cold cell-Matrigel mixture was loaded into the bottom channel of the chips followed by 20 min incubation in the incubator. After Matrigel gelation, 50 µl LUHMES pre-differentiation medium was given to the in/outlets of the bottom channel and then the top channel of the chips was coated with PBS diluted collagen-Ⅳ (100 µg/ml) and fibronectin (50 µg/ml) for 2h in the incubator. After channel coating, 50 µl LUHMES pre-differentiation medium was given to the in/outlets of the top channel and the plate was placed in the incubator, either under static condition or on the periodical perfusion rocker (MIMETAS) with the setting of 7 ° inclination and 8-min rocking interval. Specifically, the statically cultured plate was manually perfused on the rocker for 30 min daily to promote medium diffusion into the gel. Four days after neural seeding, the medium was switched to LUHMES final differentiation medium on both sides of chips. Eight days after neural seeding, endothelial cells were seeded in the top channel. Single hBMECs were prepared and resuspend in MV-2 medium at density of 20,000 cells/µl. Medium of the in/out of the top channel was aspirated and hBMECs was loaded into the channel by adding 3 µl cell solution as a drop at the inlet while aspirating from the outlet. The inlet was then supplied with 50 µl MV-2 medium and the plate was placed on its side in the incubator for 1 h to allow cell sediment against collagen gel. After hBMECs attachment, another 50 µl MV-2 medium was given to the outlet of top channel and plate was put back on the rocker in the incubator. A confluent tube should form in approximately 3 days, by which time lower serum containing MV-2 medium (2% FBS) was used in the top channel. Medium was completely changed for both sides of the chips every two days since the initial neural cell seeding. Assays were performed from day 15–16.

### Collagen genipin crosslinking

To specifically crosslink the collagen gel in the middle channel of the chips, genipin (G4796, Sigma-Aldrich) was used. Briefly, after collagen gelation, 10 mM genipin PBS solution was given to the top channel (20 µl in both in/outlet). The channel was perfused by placing the plate on the rocker in the cell incubator for 1h. After gel crosslinking, the genipin solution was aspirated and the top channel was perfused with PBS (50 µl in both in/outlet) in the incubator for 3 times, each taking 30 min except for the last one which took overnight. After washing, tri-seeding of the NVU can be followed as described previously.

### Assessment of medium diffusion in OrganoPlate model

Cell medium containing 0.5 mg/ml 70 kDa TRITC-dextran (T1162, Sigma-Aldrich) was used to access medium diffusion in the tri-culture NVU model. Dextran medium was given either in top channel to access medium diffusion from tube lumen to bottom neural cells or given in bottom channel to access medium diffusion the other way around. The plate was perfused on the rocker in the incubator for 24 h. Images were taken with fluorescence microscope to access penetration of dextran in the chips. ImageJ software was used to visualize the medium gradient.

### Calcium imaging

The Fluo-4 AM (ab241082, Abcam) DMSO stock was diluted with Neurobasal medium to make final concentration of 5 µM. Medium was aspirated from the chips and 50 µl Fluo-4 AM dye was given to the in/outlets of both the top and bottom channel. The plate was put on the rocker in the incubator for 1 h after which the chips were washed three times with medium and new medium was given. To detect spontaneous calcium fluctuations of the neurons, the chips were imaged with fluorescence microscope (20 × magnification, Ex/Em 494/506 nm) at 2 Hz for 90 seconds. Fluorescence intensity of region of interest (ROI) was measured using ImageJ software and the profile was plotted [50].

### Barrier integrity assay (BIA)

All in/outlets of the chips were first rinsed with cell medium to ensure proper hydrostatic balance at the middle collagen gel-endothelial tube surface. A total of 100 µl of endothelial medium containing 0.5 mg/ml 20 kDa FITC-dextran (FD20S, Sigma-Aldrich) was given to the top channel which contained endothelial tube (50 µl in both in/outlet). Dextran free medium was given to the bottom channel (50 µl in both in/outlet). The plate was perfused on the rocker for 30 min in the incubator. Fluorescent images of the chips were taken with confocal microscope (Yokogawa). Tube leakage index was calculated as follows: [fluorescent signal from the collagen gel (Fluo_Gel_)/fluorescent signal in tube lumen (Fluo_Lum_)].

### P-glycoprotein functionality

P-glycoprotein (P-gp) assays were adapted from reported study [51]. Briefly, mono-cultured endothelial tubes in the 3-lane chips were pre-perfused with 20 µM verapamil (BP720, Sigma-Aldrich) for 30 min followed by another 1 h perfusion of 10 µM verapamil and 1 µM calcein AM (C1430, Thermo Fisher) in the tubes. As control tubes, equal molar of DMSO was used instead of verapamil. Tubes were then washed twice with medium followed by live staining with Hoechst (#62249, Thermo Fisher, 1:1000) to stain the endothelial nucleus. Tubes images were taken by confocal microscope. Intracellular calcein intensity of endothelial cells laying at the gel interface was measured and normalized to cell counts using ImageJ software.

### Immunofluorescence microscopy (IFM)

The IFM method was adapted from the original protocol provided by the MIMETAS company. Briefly, cell culture in 3-lane chips were fixed with 4% (v/v) paraformaldehyde (PFA) (P6148, Sigma-Aldrich) for 30 min followed by 3 times washing (30 min per wash) in PBS. Cells were permeabilized with 0.3% (v/v) Triton X-100 (T8787, Sigma-Aldrich) for 30 min followed by 3 times washing (30 min per wash) in PBS. Cells were then blocked in PBS containing 2% FBS, 2% (w/v) bovine serum albumin (BSA, A2153, Sigma-Aldrich), 0.1% (v/v) Tween-20 (P9416, Sigma-Aldrich) for 3 h. Primary antibodies against VE-cadherin (vascular endothelial cadherin, #33168, Abcam, 1:400), Claudin-5 (#4C3C2, Thermo Fisher, 1:50), ZO-1 (Zonula Occludens Protein 1, #61-7300, Thermo Fisher, 1:200), S100β (S100 calcium-binding protein B, GA504, Dako, 1:400), GFAP (glial fibrillary acidic protein, #76214, Dako, 1:200), VIM (vimentin, #45939, Abcam, 1:600), β-Ⅲ tubulin (#32-2600, Thermo Fisher, 1:800), MAP-2 (microtubule associated protein 2, m2323, Sigma-Aldrich, 1:500), TH (tyrosine hydroxylase, #6211, Abcam, 1:1000) were diluted in blocking buffer supplemented with additional 1% Triton X-100. The chips were perfused on the rocker with primary antibodies overnight at room temperature, where the top channel was only given with antibodies (15 µl in both in/outlet) specifically label endothelial cells and bottom channel was only given with antibodies (25 µl in both in/outlet) specifically label neural cells. The chips were washed 3 times (30 min per wash) in PBS followed by secondary antibody incubation. Secondary antibodies including Goat anti-Rabbit Alexa Fluor-488 (A-11008, Thermo Fisher, 1:500), Goat anti-Rabbit Alexa Fluor-647 (A-21245, Thermo Fisher, 1:500), Goat anti-Mouse Alexa Fluor-568 (A-21422, Thermo Fisher, 1:500), Alexa Fluor-488 Phalloidin (A12379, Thermo Fisher, 1:40) were diluted in blocking buffer supplemented with additional 1% Triton X-100. The chips were perfused on the rocker with secondary antibodies overnight at room temperature. The chips were perfused one more hour with Hoechst (1:1000) in PBS to stain the nucleus. The chips were washed 4 times (30 min per wash) in PBS on the rocker and images were acquired by confocal microscope. For quantification, images were taken as maximum Z-projection with 20 × objective at 2 µm slicing interval and the images were analyzed using ImageJ software. For 3D image reconstruction, images were taken at 20 × magnification across whole tube structure with 2 µm slicing interval and 3D images were generated using ImageJ software.

### Tube inflammation response and PBMC interaction

Medium containing TNFα (#300-01A, Peprotech) and IL-1β (#200-01B, Peprotech) (both at 20 ng/ml) was given either in the tube lumen or in the bottom channel in/outlet of the chips followed by placing on the rocker for 24 h in the incubator. BIA was performed to check cytokines induced barrier disruption whereas IFM was performed to validate these effects from the molecular level. To check endothelial recruitment of the immune cells 24 h after exposure of TNFα and IL-1β, 100 µl freshly isolated PBMCs at the density of 2,000,000 cells/ml and pre-labeled with green CMFDA dye (C2925, Thermo Fisher), were perfused through the tube lumen at the rocking interval of 8 min/cycle. Arrest of the cells in the tubes was checked in 1 h whereas BBB extravasation was checked in 48 h. Fluorescence images were taken and migrating distance of the cells in the gel was measured using ImageJ software.

### Transwell NVU model

For the transwell-based tri-culture NVU model, the backside of the transwell insert (0.4 µm pore size) (#140652, Thermo Fisher) was coated with Matrigel overnight in the incubator followed by seeding astrocytes at 150,000 cells/cm^2^. The insert was placed inverted in the incubator to allow for astrocyte sedimentation to the backside of the insert. Meanwhile, the transwell bottom was coated overnight with PLO and fibronectin. Next day, inside of the insert was coated with Collagen-IV (50 µg/ml) and Fibronectin (25 µg/ml) for 4 h in the incubator followed by seeding hBMECs at 220,000 cells/cm^2^. Transwell bottom was seeded with 2-day pre-differentiated LUHMES cells at 100,000 cells/cm^2^. MV-2 medium was used for hBMECs in the insert (apical side) whereas 1:1 mixture of MV-2 and LUHMES differentiation medium was used for LUHMES neurons and astrocytes in the outer well (basolateral side). Medium was refreshed every two days. The tri-culture BBB model was cultured for 10 days before the assays.

### Nanobody transcytosis in transwell NVU model

Three Nbs (F12, H7, R2) were produced in house, where F12 and H7 were raised against proHB-EGF **(manuscript under review)**; R2 was raised against an irrelevant target copper containing azo-dye Reactive Red 6 [52, 53]. All Nbs were conjugated to Alexa Fluor-647 fluorophore via the C-terminal single cysteine as previously described **(manuscript under review)**. Before transcytosis assay, the inserts were moved to new wells and were treated with 0.5% FBS containing MV-2 medium (serum starvation) plus 10 µM forskolin for 2 h to transiently improve the barrier impermeability. All Nbs were prepared at 1000 nM in serum starvation medium. Nb transcytosis assay took 3 h, where at different time points (1, 2, 3 h) a portion of medium sample from basal side was collected followed by replenishing fresh medium. To detect Nb transcytosis, medium samples were subjected to SDS-PAGE followed by gel imaged by ChemiDoc MP imaging system (BioRad) using 647 nm laser line. In parallel, serial diluted samples of each Nb were run in SDS-PAGE gel to generate standard curve. Band intensities of the Nbs were measured using ImageJ software and then corrected with labeling efficiency.

### Nanobody transcytosis in OrganoPlate NVU model

All in/outlets of the 3-lane chip were first rinsed with cell medium to ensure proper hydrostatic pressure balance at the middle collagen gel-endothelial tube surface. Before the assay, the endothelial tubes were perfused with 0.5% FBS containing MV-2 medium plus 10 µM forskolin for 2 h. All Alexa Fluor-647 labeled Nbs (F12, H7, R2) were prepared at 1000 nM in serum starvation medium supplemented with 10,000 nM 20 kDa FITC-dextran. A total of 100 µl Nb medium was given to the top channel (50 µl in both in/outlet) where the endothelial tube resided. The plate was placed on the rocker with the rocking interval of 2 min/cycle and incubated for 3 h. At the 30 min timepoint all chips were first checked with BIA to ensure good quality of the endothelial barrier integrity where tubes with leakage index above 15% were discontinued. After the assay, endothelial tubes were rinsed once with cell medium, and the chips were immediately fixed with 4% PFA and IFM was followed to determine Nb transcytosis.

Apart from single Nb administration, a dual-Nb transcytosis setting was also explored in the NVU model. Briefly, Alexa Fluor-647 labeled Nbs (F12, H7, R2) was mixed with equimolar Alexa Fluor-488 labeled R2 at the concentration of 500 nM each. Tubes were perfused with the 1000 nM Nb mixture and transcytosis was measured via IFM as described above.

### Statistical analysis

All data were presented as mean ± standard deviation (SD). Statistical analysis was performed by student’s t-test or one way ANOVA method using GraphPad Prism 9 software. For multiple comparisons, either Tukey’s multiple comparisons tests or Dunnett’s multiple comparisons tests was performed. P values < 0.05 were considered statistically significant.

## Results

### Screening for optimal 3D NVU model design

Each chip of the 3-lane OrganoPlate contains three adjacent channels that are separated by a membrane-free structure called Phaseguide **(Fig. 1a)** [54]. The collagen-Ⅰ ECM gel was loaded in the middle channel of the chips whereby it supported the adhesion and growth of the endothelial cells in the top channel. Anatomically, as brain capillaries are closely surrounded by the astrocyte endfeet, we first thought to seed astrocytes directly in the collagen gel and expected a close contact to form between the astrocytes and the endothelial tube. As a result, astrocytes were found to nicely survive in the collagen gel and could form glia-endothelial interactions **(Fig. S1a)**. However, these expanding and stretching astrocytes profoundly distorted the gel over time and eventually made the whole model unsustainable, which was in line with previous reports [30, 42]. Besides, we found that neurons were unable to survive in the collagen gel **(data not shown)**. To tackle this, we introduced a double-gel design in the chip, where the bottom channel was additionally filled with Matrigel to provide a 3D culture environment for the neural cells **(Fig. 1b)**. As a result, growing astrocytes in the bottom Matrigel avoided obvious disruptions to the collagen gel while still allowing the cells to migrate through the gel to contact the endothelial tube **(Fig. S1b)**. Furthermore, neurons growing in the Matrigel were able to differentiate and form neural networks **(Fig. 2 and Fig. 3a)**. Importantly, the presence of collagen gel was found to totally retain the neuron somas in the Matrigel meanwhile allowing for efficient neurite outgrowth in the collagen **(Fig. S2)**.

**Fig. 1.**
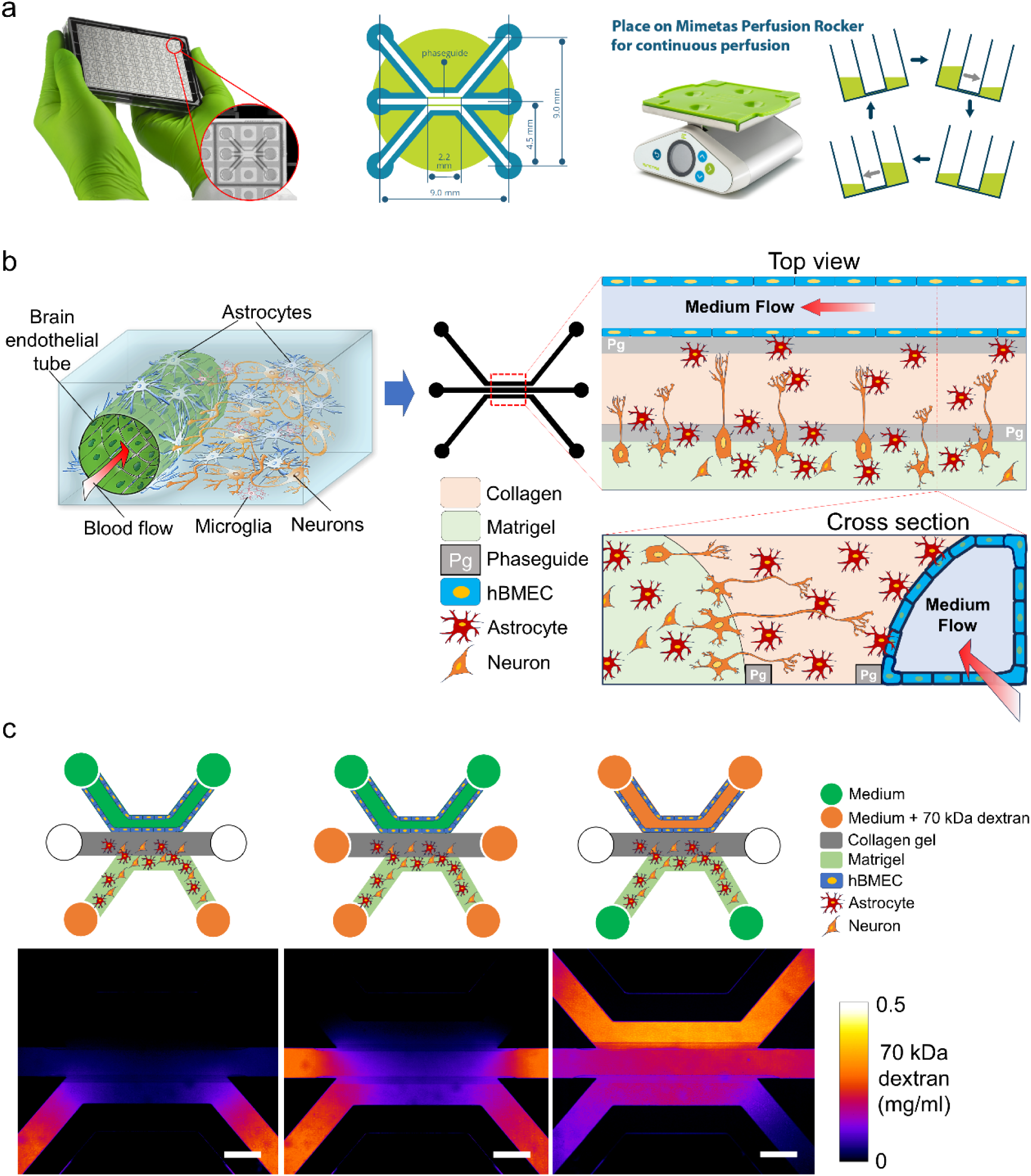
Design of a full-3D NVU model in the 3-lane OrganoPlate chips. **(a)** Schematics of the 3-lane chip. The copyright images are adopted with permission from MIMETAS BV. **(b)** Schematic illustration of a simplified NVU and its fulfillment in the 3-lane chip. **(c)** Medium diffusion pattern in the double-gel designed NVU model. Medium supplemented with fluorescently labeled dextran is given to different in/outlets of the model to allow for 24 h incubation. Images were taken as fluorescence. Scale bars are 500 µm.

**Fig. 2.**
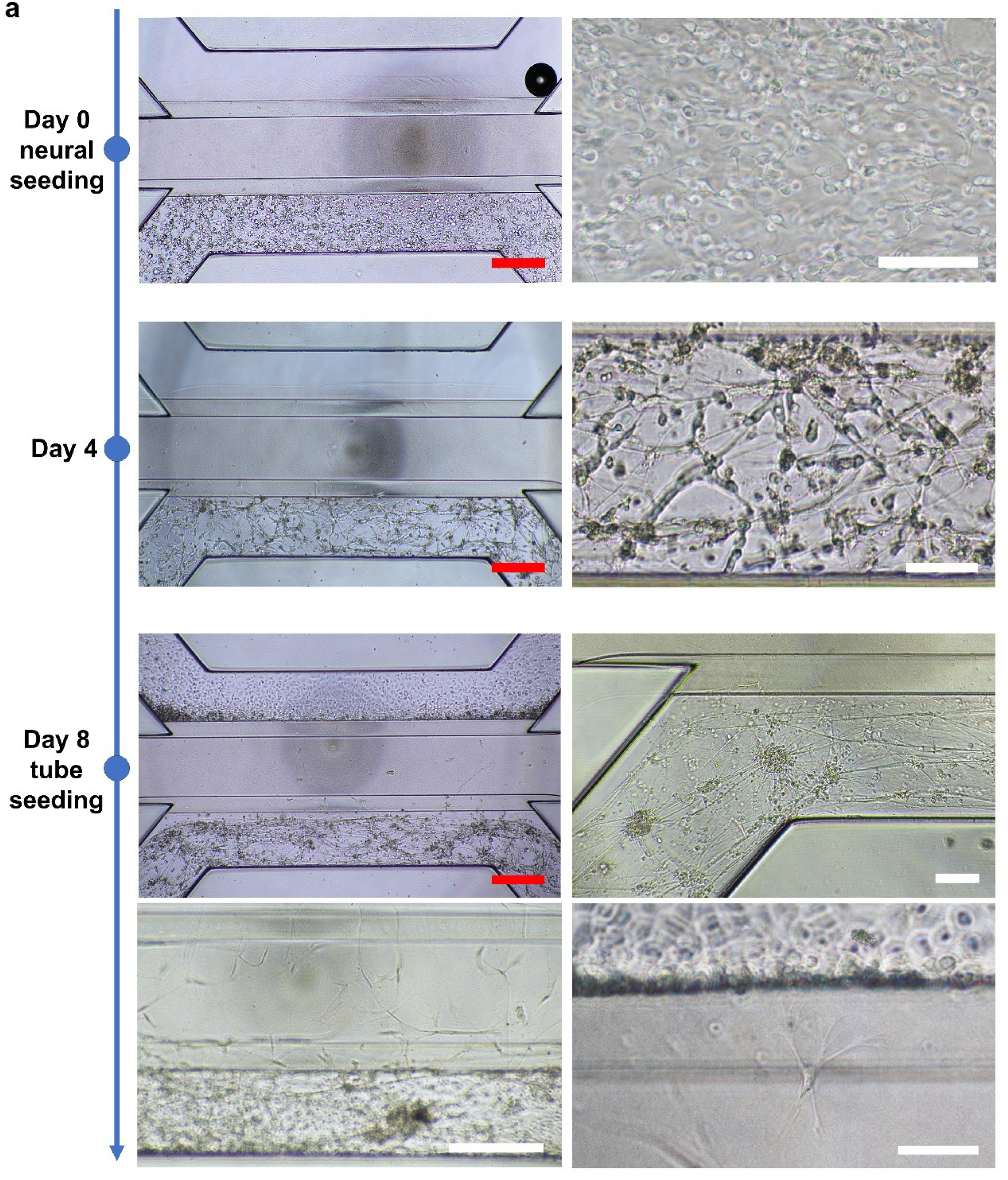
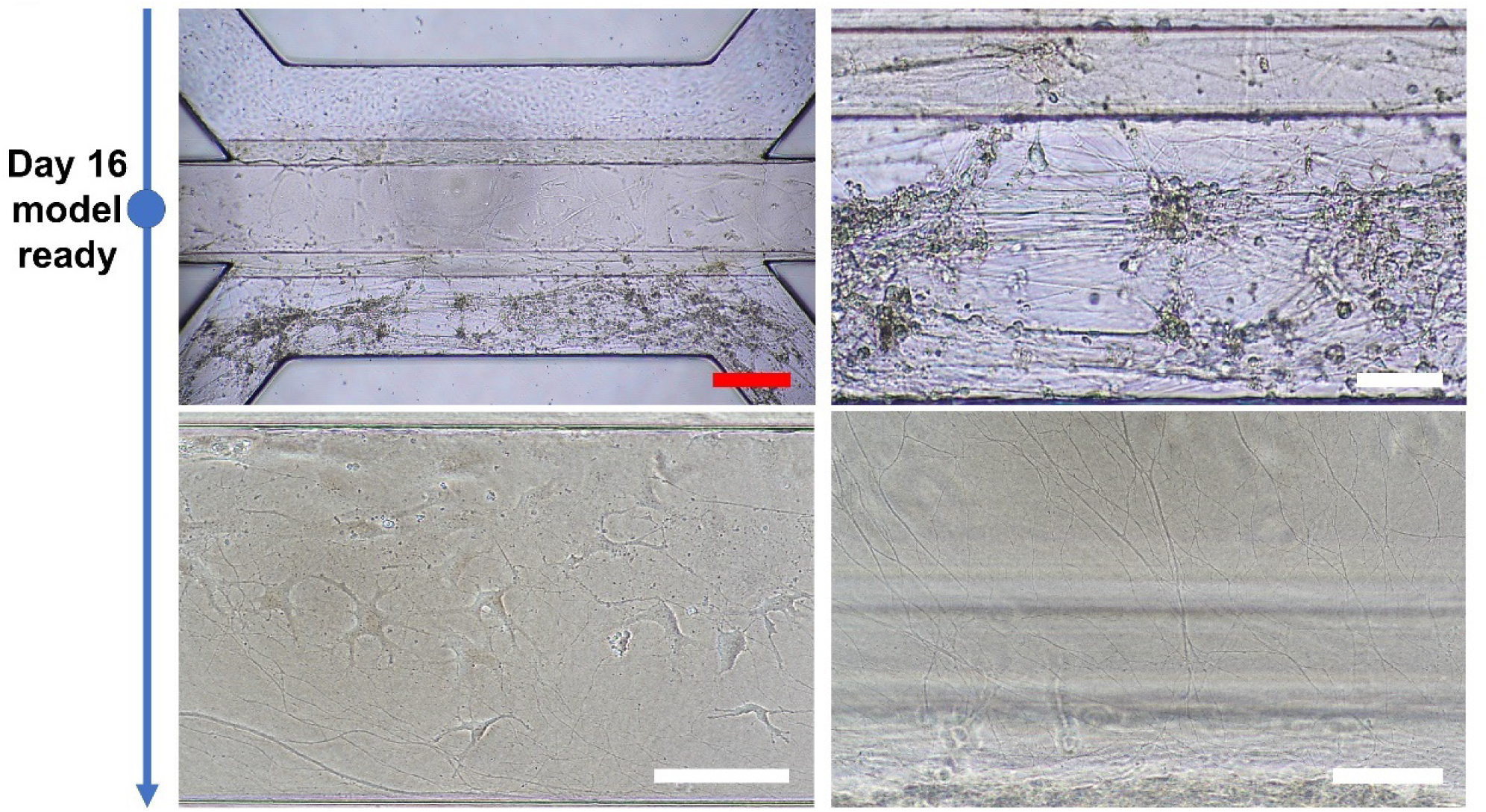
Generation of a full-3D NVU model in the 3-lane OrganoPlate chips. Timeframe images of the NVU modeling protocol. Images were taken as bright field. Scale bars are 500 µm (red bars) or 100 µm (white bars).

**Fig. 3.**
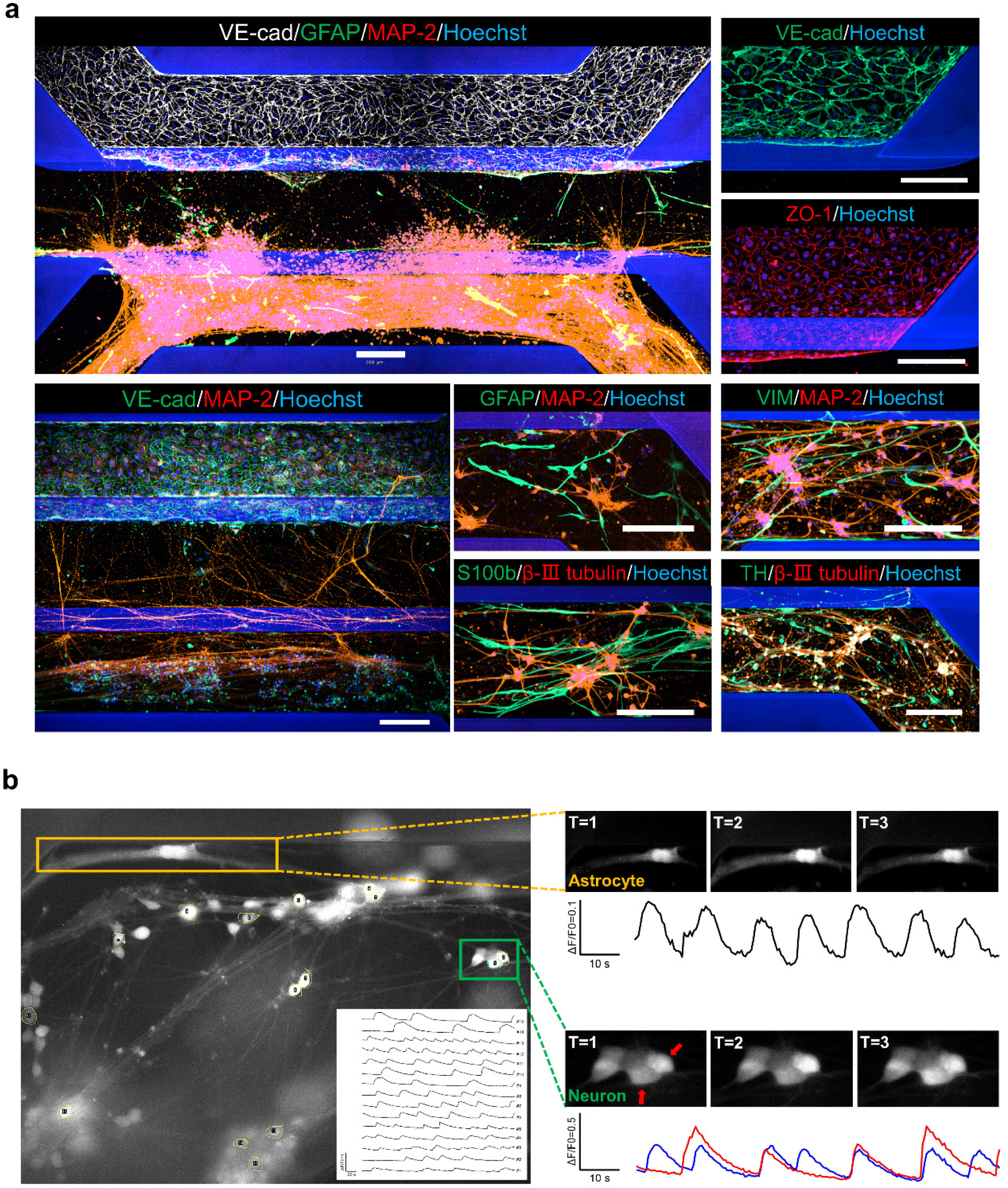
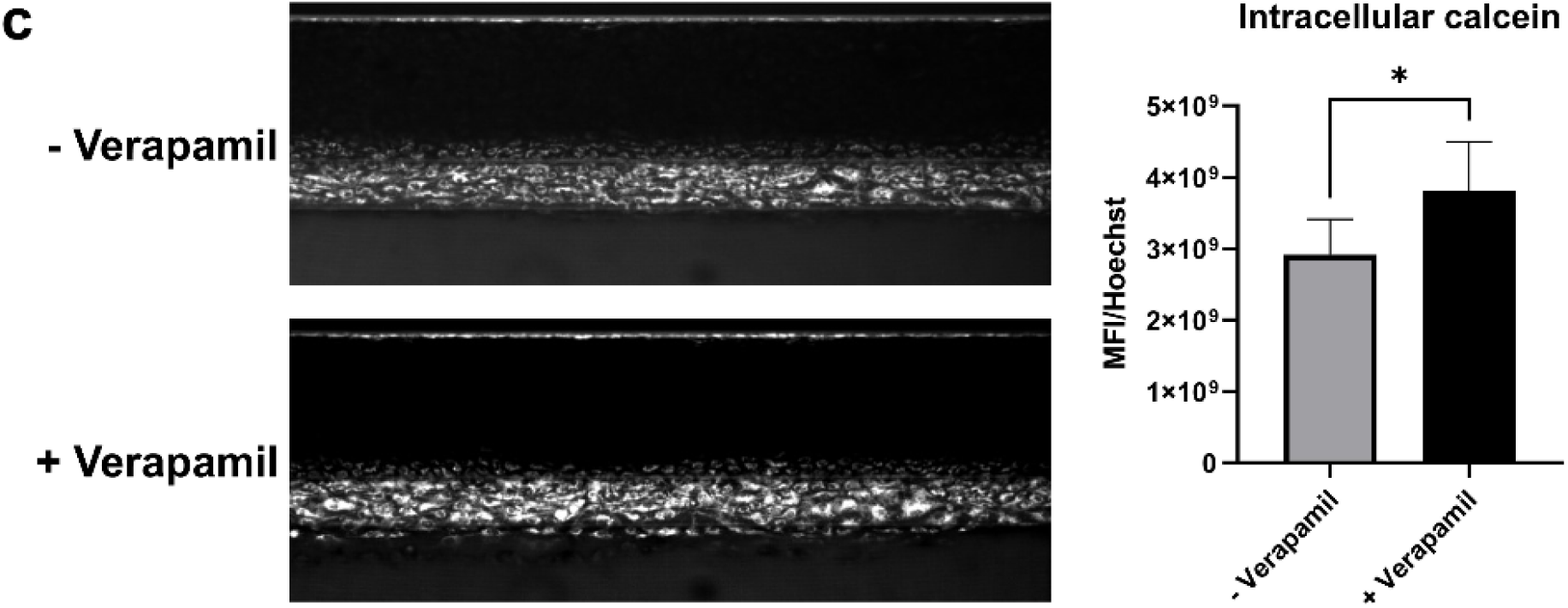
Characteristics of the NVU model. **(a)** IFM images showing the expressions of the endothelial-, astrocyte- and neuron-specific markers in the NVU model. All IFM images were taken at maximum projection mode. Scale bars are 200 µm. **(b)** Calcium imaging showing spontaneous activities in the neural cells. Inset showing 15 randomly selected neurons in the field. Zoom in image (orange) showing spontaneous activities in an astrocyte. Zoom in image (green) showing signal transmission among neurons within a neuron cluster. Calcium signals were acquired at 2 Hz. **(c)** P-gp assay showing efflux function of the endothelial tubes. Images were taken as fluorescence. For quantification, each group contains 5 chips. Graph shows mean ± SD. Statistical analysis was performed using student’s t-test; *P < 0.05.

For multicellular BBB or NVU modeling in the chips, the bottom channel is normally filled with neural medium [46, 47]. However, as we loaded the bottom channel with ECM gel, sufficient medium supply of the neural cells remains a challenge. To ask if neural cells in deeper gel area (i.e., observation window area of the chip) can still be properly fed, we evaluated the medium diffusion pattern in the model by supplying 70 kDa TRITC-dextran in the medium. Despite at a slow speed, it was confirmed that large molecules (∼70 kDa) added in the bottom channel in/outlets are able to diffuse and infiltrate the double-gel layer within 24 h **(Fig. 1c left)**. Due to presence of the tube and hydrostatic pressure in the top channel, neural medium totally stopped behind the endothelial wall. In addition, we found that giving extra neural medium to the in/outlets of middle channel did not substantially improve its infiltration to the center gel area within 24 h **(Fig. 1c middle)**. Moreover, this was shown to induce angiogenesis of the tube **(Fig. S3)**. We also checked the diffusion pattern of the endothelial medium into the gel. It was found that dextran in the tube lumen infiltrated the whole chip in 24 h, which could be a result of both the active intracellular transport and paracellular leakage of the endothelial barrier over such a long time **(Fig. 1c right)**. In line with other NVU models, these results suggested that the neural cells in the gel were actually facing a hybrid culture environment consisting of medium from both the “blood side” and the “brain side” [30]. This is also a true reflection of the *in-vivo* situation that the BBB is slightly permeable to large blood-derived molecules, such as albumin [55, 56]. Taken together, we showed that the double-gel design is suitable for modeling NVU in the 3- lane OrganoPlate.

### Characteristics of the full-3D NVU model

To give neurons enough time to differentiate, our protocol was segregated by a neural monoculture phase and an endothelial co-culture phase **(Fig. 2)**. After collagen loading in the middle channel, Matrigel encapsulating astrocytes and pre-differentiated LUHMES neurons was dispensed into the bottom channel and cultured alone for the first 8 days. The neurons developed networks in the bottom channel over time, meanwhile neurite outgrowth as well as astrocyte migration in the collagen gel started to be seen. It remains unclear whether the bi-directional medium flow, induced by periodical rocking of the MIMETAS plate rocker, interferes with neuron differentiation and neural network formation. To investigate this, we cultured the day-zero neurons in the chips with or without active medium perfusion. Morphologically, we did not see substantial changes during neurite outgrowth **(Fig. S4)**. Neurons under both conditions were able to develop their networks to a similar complexity. On day 8, endothelial cells were seeded in the top channel and the plates were subjected to periodical rocking to promote tube formation. An intact tube formed in 3 days and was cultured up to 9 days during the experiment. During this period, neurite outgrowth and astrocyte migration in the collagen gel channel began to accelerate. By day 16, astrocytes were in close contact with the endothelial tube, indicative of glia-endothelial communications **(Fig. 2a and Fig. 3a)**. Neurons in the Matrigel channel formed clusters and thick nerve bundles within the networks. Notably, their axons have grown through the collagen gel and formed extensive neural connections with the endothelial tube **(Fig. 2a and Fig. 3a)**. Via IFM, the endothelial tube was outlined by intercellular VE-cadherin and ZO-1. The LUHMES neurons were positive for β-Ⅲ tubulin, MAP-2 and TH. The Astrocytes showed expression of S100β, GFAP and VIM (Fig. 3a). The NVU model collapsed after prolonged culture (e.g., more than 21 days), as depicted by the shrinking of the double-gel structure and the un-sustained tube morphology **(Fig. S5)**. Nonetheless, such a long culture period granted the neurons enough time to form complex ganglia-like structures.

Neural activities in the model were assessed by calcium imaging. As was seen, both the neurons and astrocytes exhibited spontaneous calcium waves **(Fig. 3b)**. Moreover, synchronized calcium signals were noticed among the cells, indicating neural network formation. Even though the endothelial cells are cultured in a tube format, only those growing at the tube-collagen interface are physiologically polarized and are therefore most relevant for BBB transport or efflux studies in the model. P-glycoprotein (P-gp) is one of the efflux transporters on the BBB which pumps a range of drugs out of the brain [57]. For efflux assay, we perfused calcein AM, a substrate for P-gp, in the tube lumen. After entering the endothelial cells, calcein AM was metabolized into calcein which emitted fluorescence. In the presence of verapamil, a P-gp inhibitor, a stronger intracellular fluorescence could be detected inside the cells, indicating less calcein AM was pumped out **(Fig. 3c)**. This proved well-functioning of the P-pg of the endothelial tubes.

### Comprehensive assessment of different culture conditions on tube integrity

Intracellular transport studies require leak-tight endothelial barrier models as paracellular leakage of the endothelium could lead to false positive results. Tube tightness in the OrganoPlate is easily accessed by the BIA **(Fig. 4a)**. Many groups have screened for culture conditions that could yield optimal tube impermeability, such as medium choices [46] or culture additives [58, 59]. Basically, the tubes were cultured in the commercial MV-2 medium as we previously found this medium elicited an optimal barrier integrity in a transwell BBB model **(manuscript under review)**. First, we performed BIA at different time points of the mono-tube cultures to determine the window for an optimal barrier integrity. The tubes cultured for 5 to 7 days showed relatively less leakage to 20 KDa dextran whereas after 7 days they gradually lost barrier integrity **(Fig. 4b)**. In line with an early study, this indicated a loss in the tube quality after prolonged *in-vitro* culture [46].

**Fig. 4.**
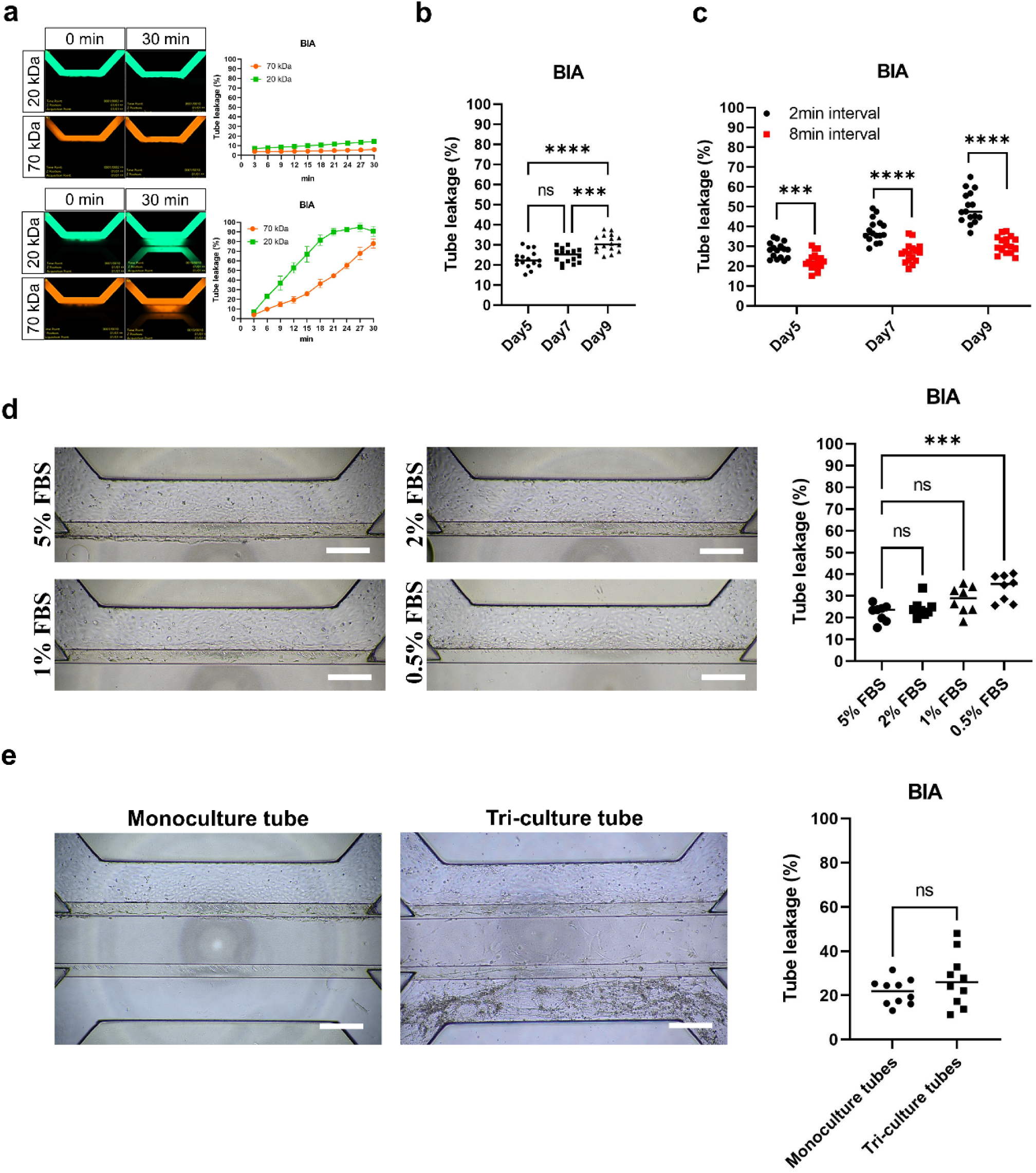
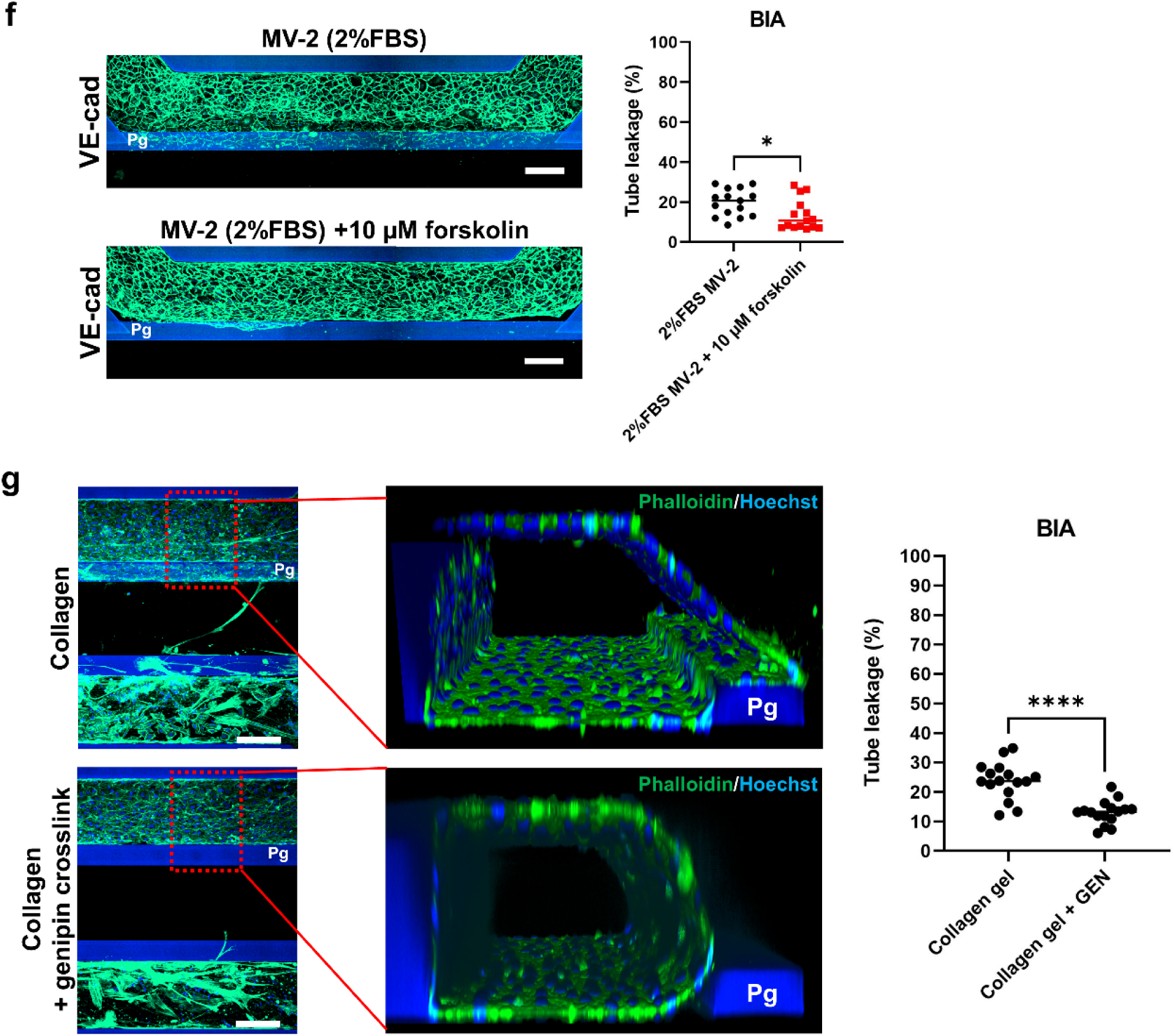
Tube integrity under different culture conditions in the NVU model. **(a)** Illustration of the BIA. Tube lumen is perfused with fluorescently labeled dextran for 30 min. Fluorescence leakage from tube lumen (Fluo_Lum_) into the adjacent collagen gel (Fluo_Gel_) is imaged over time. The ratio Fluo_Gel_/Fluo_Lum_ indicates the tube leakage levels. Examples of both the leak-tight and the leaky tubes are presented. **(b)** Time-course BIA of the mono-tubes. For quantification, each group contains 16 chips. **(c)** Effects of different plate rocking intervals on tube integrity. For quantification, each group contains 16 chips. **(d)** Effects of different FBS levels on the morphology and integrity of the mono-tubes. Images were taken as bright field. For quantification, each group contains 8 chips. **(e)** Effect of neural co-culture (astrocytes and neurons) on tube integrity. Images were taken as bright field. For quantification, each group contains 10 chips. **(f)** IFM images showing the effect of forskolin on the morphology and integrity of the mono-tubes. For quantification, each group contains 10 chips. **(g)** IFM images showing the effect of collagen genipin crosslinking on the morphology and integrity of the co-cultured tubes. For quantification, each group contains 16 chips. Scale bars are 500 µm for **panel d, e** and 200 µm for **panel f, g**. Pg: Phaseguide. All graphs show mean ± SD. Statistical analysis was performed using one-way ANOVA followed by Tukey’s multiple comparisons tests for **panel b** and Dunnett’s multiple comparisons tests for **panel d**, and student’s t-test for **panel c, e, f, and g**; ns indicates no significant differences; *P < 0.05, **P < 0.01, ***P < 0.001, ****P < 0.0001.

The flow-derived shear stress is a key contributor to tube integrity [21, 24, 26]. The MIMETAS rocker uses the shaking setting of a 7° inclination and an 8-min interval to periodically generate the bi-directional flow in the chip. However, under this setting, medium flow exists for as short as approximately 2 min per cycle. Thus, we asked if a continuous existence of the flow, achieved by setting the rock interval at 2-min, could give better tube integrity. On the contrary, tubes under such frequent bi-directional flow developed worse barrier tightness over time as compared to the tubes with 8-min rocking interval **(Fig. 4c)**.

Our early work proved the importance of serum in maintaining optimal endothelial barrier integrity **(manuscript under review)**. In the OrganoPlate, we investigated how different serum levels affected the tube integrity. Compared to 5 % serum that was originally provided by the MV-2 medium, tube tightness gradually decreased as serum level dropped **(Fig. 4d)**. In parallel, tubes cultured with lower serum showed less cell confluency and flawed tubular morphology, which was in line with other study [60]. Notably, despite that 5 % serum yielded relatively less tube leakage, it caused overgrowth of the endothelial cells, as indicated by the observation that the tubes expanded towards the middle channel at a higher speed **(Fig. 4d)**. Moreover, along with tube expansion, the Phaseguide structure of the chip began to reveal a rectangular surface for the endothelial cells to grow on **(Fig. 4g)**, which was commonly seen in many OrganoPlate-based tubular models [46, 47, 51, 61, 62]. This led to an aberrant tube morphology and could potentially compromise tube properties. Hence, as a comprehensive consideration, we chose to culture the tube with 2% serum in the endothelial medium as this serum level did not significantly disrupt barrier integrity and meanwhile could slow down tube expansion.

Co-culture with astrocytes in the BBB model has been shown to improve the properties of the endothelial barrier [39, 63–68]. In addition, neurons were found to indirectly regulate BBB integrity [39, 63, 69, 70]. In our NVU model, however, we did not see an improved barrier integrity when culturing with astrocytes and neurons **(Fig. 4e)**. In contrast, some tri-culture chips even showed worse tube tightness. Notably, we found the BIA data points of the tri-culture chips revealed a more scattered clustering, similarly to a recent study [29]. This could indicate a compromised consistency in tube quality of the tri-culture models.

Administration of cyclic adenosine monophosphate (cAMP) has been found to effectively improve barrier integrity [58, 71, 72]. We investigated the effect of forskolin, a cAMP pathway activator, on tube tightness. As a result, 10 µM forskolin was shown to significantly improve barrier resistance **(Fig. 4f)**. In addition, barrier enhancement by transient serum starvation and forskolin stimulation was more pronounced **(Fig. S6)**. On the other hand, forskolin treatment was also found to contribute to good tube morphology, as was revealed by the presence of a smooth tube border and the delayed tube expansion towards the middle channel **(Fig. 4f)**. This could be explained by the cAMP signaling pathway which arrested cell cycle of the endothelial cells [73].

Stiffness of the ECM gel was found to positively correlate with endothelial integrity and tube stability [30, 74]. We implemented genipin, a non-toxic extract from gardenia fruit that was known to crosslink collagen, to increase the gel stiffness [75]. We found that the tubes were forced to stay unexpanded and maintain a smooth border at the gel-cell interface due to their inability to degrade the crosslinked collagen **(Fig. 4g)**. Importantly, these tubes exhibited significantly better integrity than those growing against normal collagen gel **(Fig. 4g)**. However, the crosslinked collagen totally separated the neural cells and the tube from physical contact, as neither neurons nor astrocytes were able to grow into it. Nonetheless, large molecules such as 150 kDa dextran were still able to freely diffuse in the crosslinked collagen gel **(data not shown)**.

### Neuroinflammation induced tube disruption

Inflammatory cues released by the reactive glial cells are major sources of neuroinflammation that lead to BBB damage [76–80]. Here we tested the possibility of modeling neuroinflammation-induced BBB disruption by exposing the tube model to inflammatory cytokines. TNFα and IL-1β were given to either the tube lumen (blood side) or the bottom channel (brain side) and the responses of the brain endothelial tubes were investigated 24 h later. It was found that the tubes underwent a significant cell loss after exposure to these stimuli **(Fig. 5a, b)**. The remaining endothelial cells stretched to cover the lost area, as revealed by their enlarged cell body and the reformation of junctional connections, which is similar to the response of a physically insulted endothelial layer [25]. In parallel, cell morphology changed from the tightly packed cobblestone-like shape to the sparsely distributed spindle-like shape, followed by cell alignment along the flow direction **(Fig. 5a)**. We next examined the expression of skeletal and junctional proteins of the endothelial cells after the treatment of TNFα and IL-1β. The expression of skeletal protein F-actin was found to be unaffected whereas the expression of junctional proteins VE-cadherin, ZO-1 and Claudin-5 were found to be significantly decreased and further accompanying an increased tube leakage **(Fig. 5c)**. Notably, for these abovementioned alterations of the endothelial tubes, the effects were found to be more pronounced when the cytokines were given to the tube lumen. It has been shown that endothelial damage caused by inflammatory stimulation follows a dose-dependent principle [27, 81, 82]. Thus, we speculated that the cytokines given at the bottom channel could have been diluted in the gel along their way of reaching the basal side of the tubes. As proof, we found that TNFα and IL-1β used at a higher concentration (50 ng/ml) in the bottom channel resulted in an increased tube leakage as compared to the use at 20 ng/ml **(Fig. 5d)**.

**Fig. 5.**
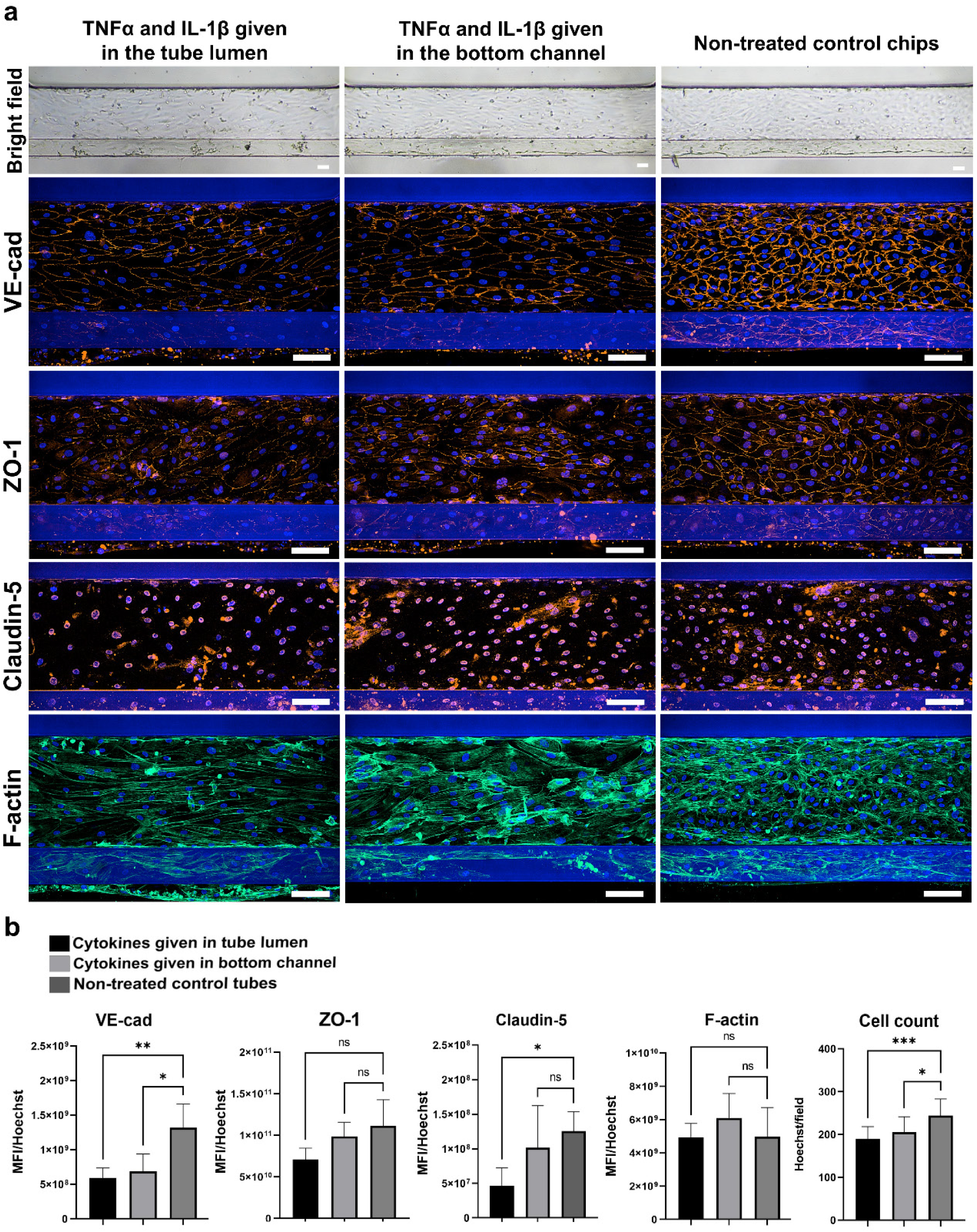
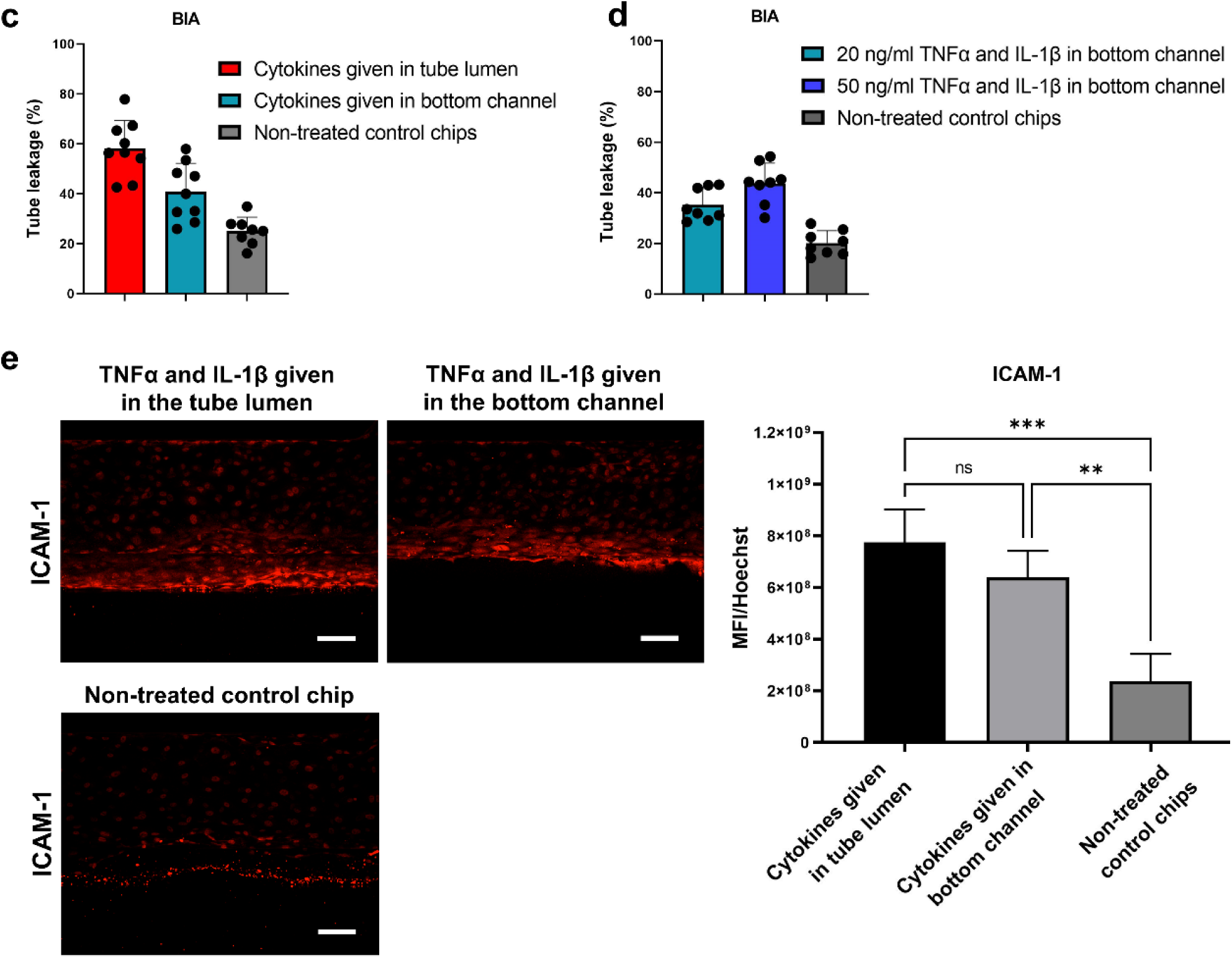
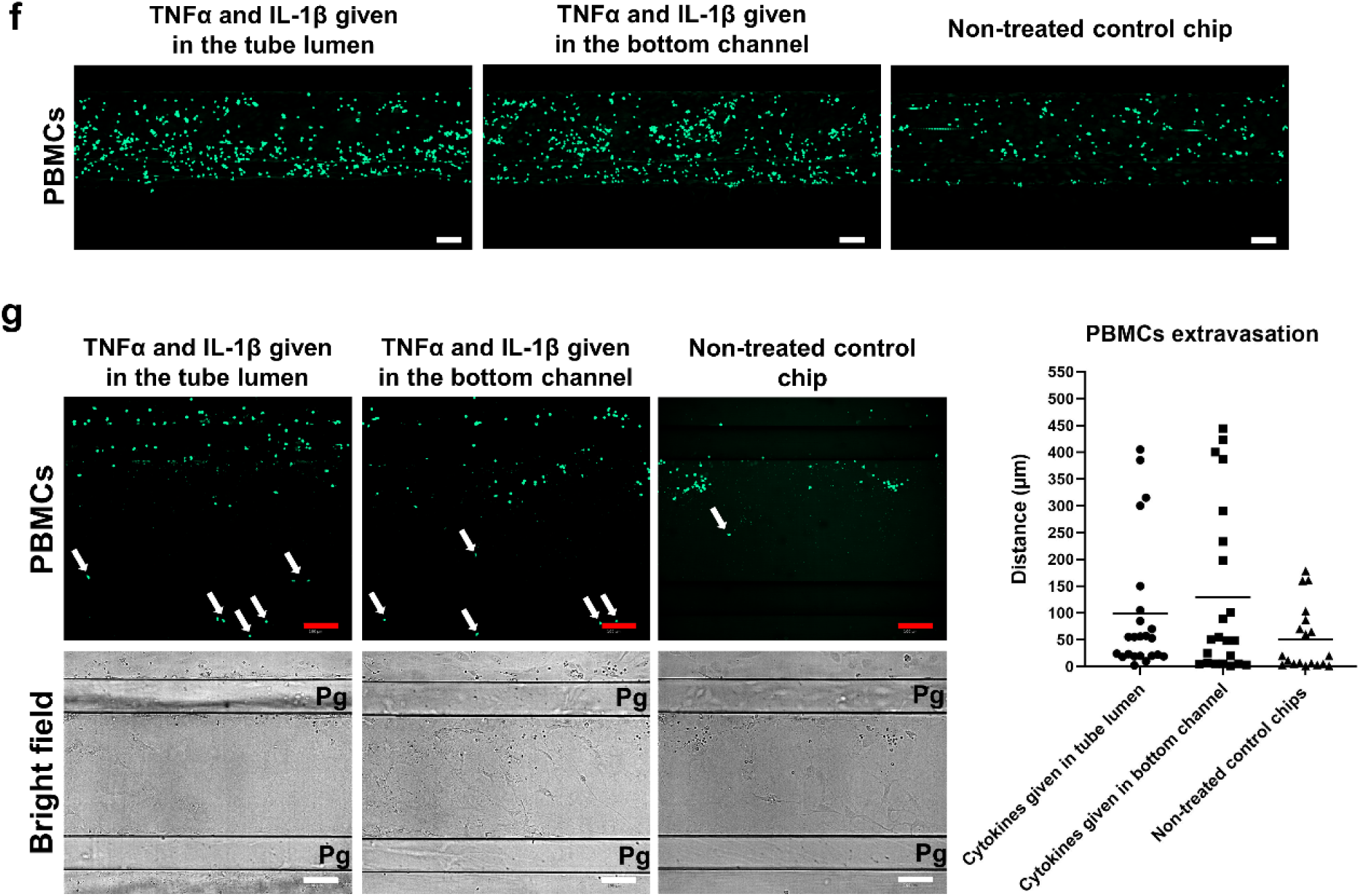
Immune responses of the brain endothelial tubes to TNFα and IL-1β stimulation in the NVU chip model. **(a)** Effects of 24 h stimulation of TNFα and IL-1β on junctional protein expressions and tube morphology. 20 ng/ml TNFα and IL-1β were given either in the tube lumen or in the bottom channel of the model. Data was acquired by IFM. **(b)** Quantification of the junctional protein expressions indicated in (a). Expression was normalized to counted cell number. Each group contains 4 chips. **(c)** Effects of 24 h stimulation of 20 ng/ml TNFα and IL-1β on tube integrity. For quantification, each group contains 8 chips. **(d)** Dose-dependent effects of TNFα and IL-1β stimulation on tube integrity when given in the bottom channel of the model. For quantification, each group contains 8 chips. **(e)** Effect of 24 h stimulation of 20 ng/ml TNFα and IL-1β on the expression of ICAM-1 in the endothelial cells. IFM images were taken to assess the expression. Expression was normalized to counted cell number. For quantification, each group contains 4 chips. **(f)** Effect of 24 h stimulation of 20 ng/ml TNFα and IL-1β on PBMCs recruitment potential of the tubes. Tubes were perfused with fluorescently labeled live PBMCs for 1h following cytokine stimulation. **(g)** Effect of 24 h stimulation of 20 ng/ml TNFα and IL-1β on BBB extravasation and migrating potentials of the PBMCs. Tubes were perfused with fluorescently labeled live PBMCs for 48 h following cytokine stimulation. Migrating distance is the length that the cells travel in the gel after leaving the tube edge. For quantification, data was pooled from 2 chips in each group. Pg: Phaseguide. All scale bars are 100 µm. All IFM images were taken at maximum projection mode. All graphs show mean ± SD. Statistical analysis was performed using one-way ANOVA followed by Tukey’s multiple comparisons tests; ns indicates no significant differences; *P < 0.05, **P < 0.01, ***P < 0.001, ****P < 0.0001.

Inflamed BBB upregulates a number of adhesion molecules which in turn efficiently recruit the circulating immune cells and favor their extravasation [83]. We found that the intercellular adhesion molecule 1 (ICAM-1) in the endothelial cells significantly elevated 24 h after the treatment of TNFα and IL-1β **(Fig. 5e)**. Tube perfusion with human PBMCs **(Fig. 5f)** revealed a significantly higher cell arrest in the inflamed tubes in as short as 1 h. For BBB extravasation, a higher number of PMBCs with a longer infiltrating distance were detected in the TNFα and IL-1β treated chips, with some PMBCs being able to penetrate the entire collagen section to reach the bottom channel **(Fig. 5g)**. There was no significant difference in the number of extravasated PBMCs or migrating distance between the chips whether the cytokines were given to the tube lumen or the bottom channel.

### Nb transcytosis in NVU model

We previously identified two proHB-EGF targeted Nbs (named F12 and H7) that are able to cross the BBB via receptor-mediated transcytosis (RMT) **(manuscript under review)**. Here, F12 and H7, as well as a negative Nb control R2, were tested in the NVU chip model, and the results were compared with that from a transwell model counterpart **(Fig. S7a)**. To rule out discrepancies resulting from other than the model format itself, both models shared the same cell types, medium compositions, and were incubated with equimolar Nbs. We first compared Nb association (i.e., binding and uptake) in the endothelial cells between the two models. In the transwell model, despite that F12 and H7 showed better endothelial uptake than the negative R2, accumulation of R2 in the endothelial cells was also broadly detected **(Fig. S7b)**, which we speculated to result from nonspecific binding of R2 to the endothelial cells owing to the static culture condition. In the chip model, the interaction of R2 to the tubes was greatly reduced whereas F12 and H7 still preserved strong endothelial association **(Fig. 6a)**. This suggested that the nonspecific weak binding of R2 to the endothelial tubes was greatly reduced under flow condition. Interestingly, we found that F12 and H7 cumulated more efficiently along the tube-collagen interphase **(Fig. 6a)**, indicating endothelial cells growing at this area were more actively and specifically interacting with the Nbs. Previously, we showed endothelial polarization (e.g., growing in the transwell system) was a necessary inducer for sufficient surface display of proHB-EGF **(manuscript under review)**. Here, the result indicated that growing against the collagen gel also induced proper polarization of the endothelial cells which then triggered the enrichment of proHB-EGF on the endothelial surface.

**Fig. 6.**
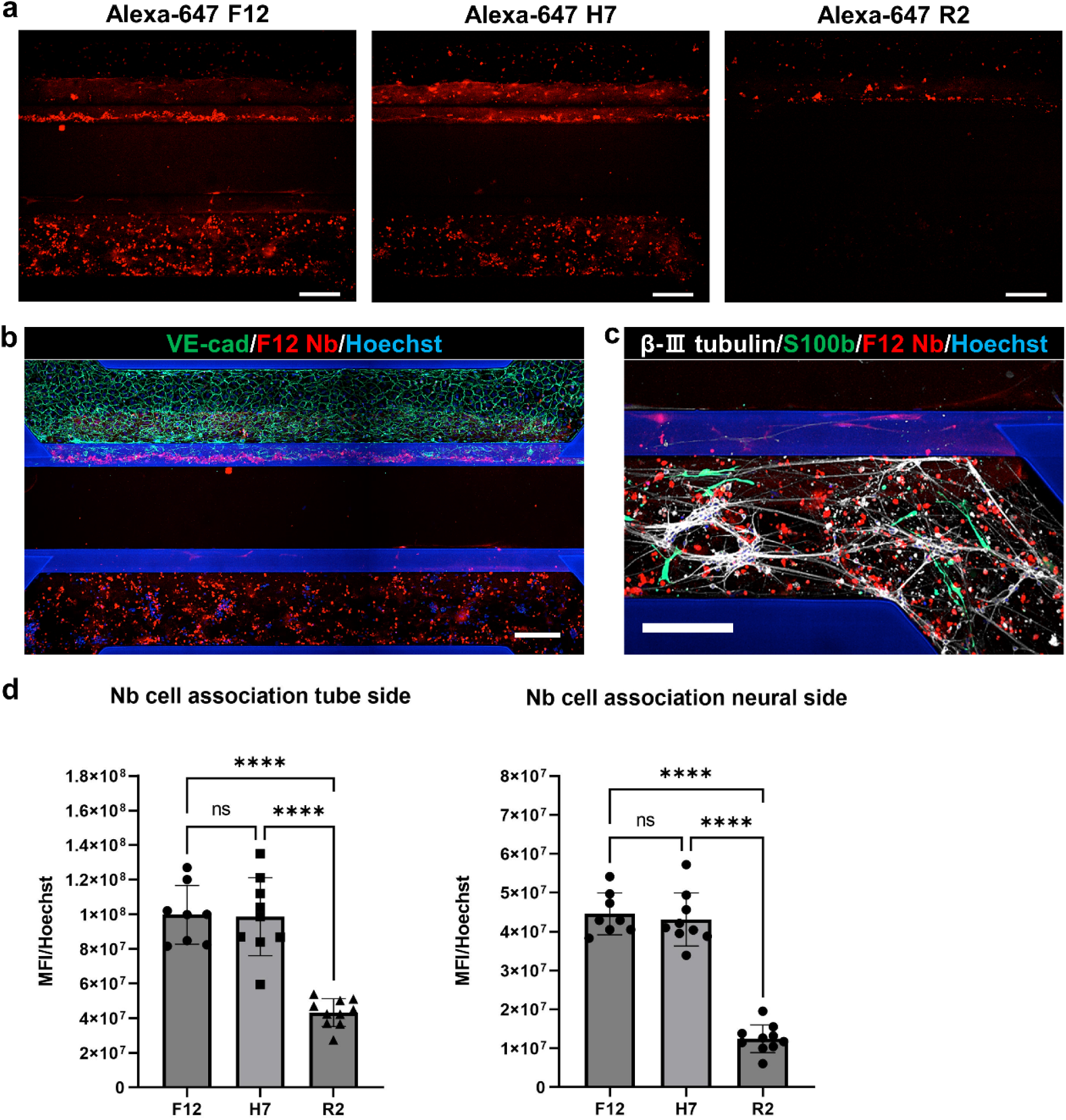
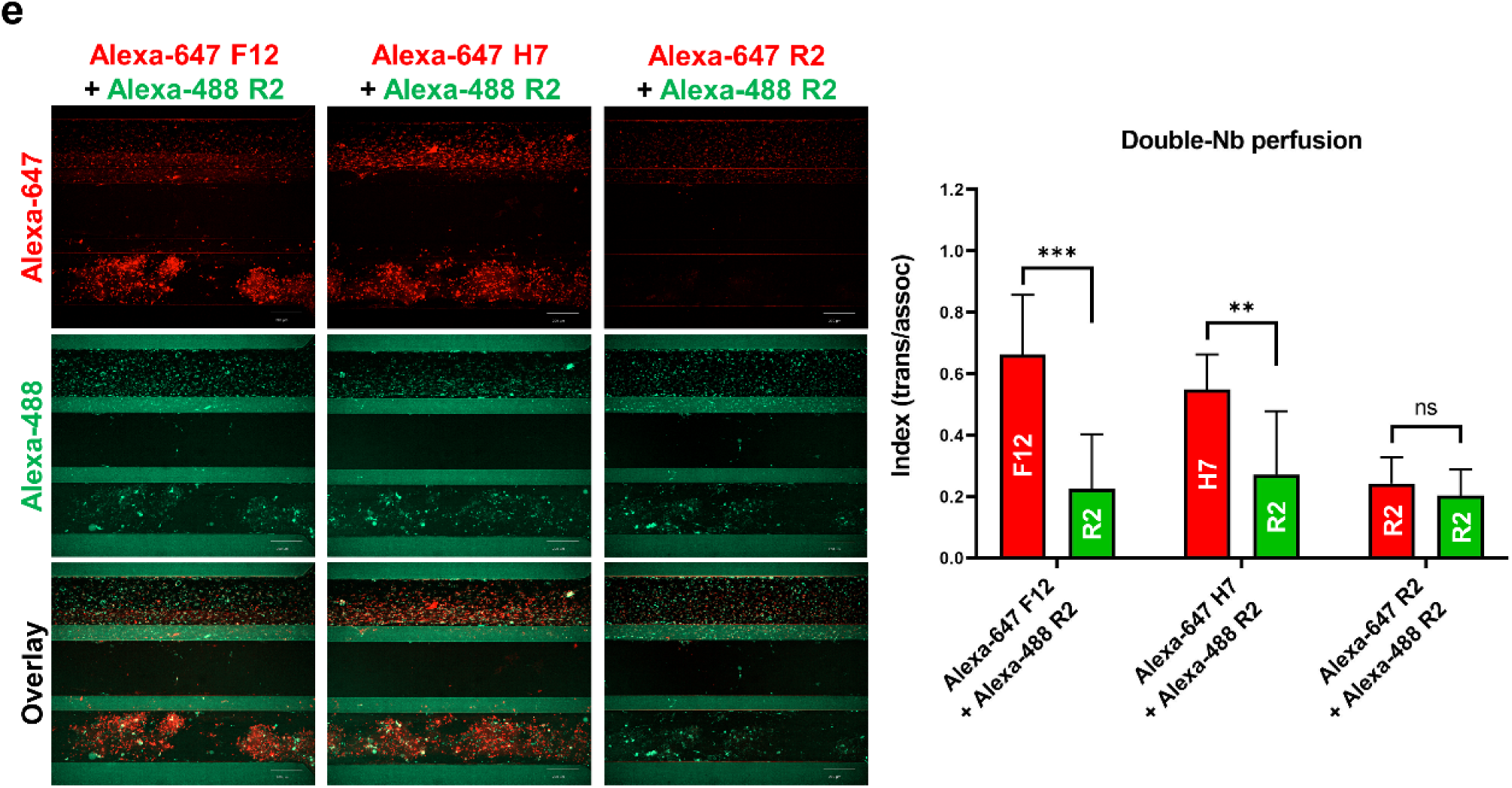
Nb transcytosis in the OrganoPlate-based NVU model. **(a)** 1000 nM Alexa Fluor-647 labeled Nbs, where F12 and H7 target proHB-EGF and R2 targets Reactive Red 6, were perfused in the tube lumen for 3 h. Fluorescence signal of the Nbs was acquired to indicate Nb transcytosis in the bottom channel. **(b)** IFM image showing the transcytosed Nbs (e.g., F12) in the bottom channel. **(c)** High magnification IFM image showing the transcytosed Nbs (e.g., F12) in the bottom channel. **(d)** Quantification of endocytosed Nbs in the endothelial cells and transcytosed Nb in the bottom channel. Each Nb group contains 8-10 chips. Fluorescence signal of the Nbs was normalized to counted cell number. **(e)** Equal molar (500 nM) of Alexa Fluor-647 labeled Nb (F12, H7, R2) and Alexa Fluor-488 labeled R2 were co-perfused in the tube lumen for 3 h. Fluorescence signal of the Nbs was acquired by confocal microscopy. The trans/assoc index was calculated for both Alexa Fluor-647 Nb (F12, H7, R2) and Alexa Fluor-488 R2. Each Nb group contains 8 chips. All scale bars are 200 µm. All IFM images were taken at maximum projection mode. All graphs show mean ± SD. Statistical analysis was performed using one-way ANOVA followed by Tukey’s multiple comparisons tests for panel d and student’s t-test for panel e; ns indicates no significant differences; *P < 0.05, **P < 0.01, ***P < 0.001, ****P < 0.0001.

Next, Nb transcytosis was investigated in the two models. In the transwell model, both F12 and H7 demonstrated significantly higher BBB transport than R2 **(Fig. S7c)**. Notably, for the chip model, as medium sampling from the bottom channel was barely feasible with our double-gel design, Nbs crossing the brain endothelial tubes cannot be accurately quantified in the same way as was done in the transwell model. Instead, we measured the Nb signal directly in the chips by fluorescence microscopy and considered the signal cumulated in the bottom channel as transcytosed Nbs **(Fig. 6b, c)**. Results showed that both F12 and H7 had significantly more depositions in the bottom channel than R2 **(Fig. 6d)**, which was similar to the extent of their cumulation in the tube lumen. For comparisons between the two models, we processed the Nb data in a fold-change manner to demonstrate the predominance of F12 and H7 in either BBB association or transcytosis when comparing with R2. In both cases, the values of F12/R2 and H7/R2 were higher in the chip model **(Table 1)**. The higher values suggested that the nature and potential of F12 and H7 in active BBB targeting and transcytosis could be better revealed by the chip model.

**Table 1.** Fold-change mannered comparison of BBB association and transcytosis of the Nbs between two models.

|  | Ratio | Transwell model | Chip model |
| --- | --- | --- | --- |
| BBB association | F12/R2 | 1.312±0.170 | 2.3866±0.6028 |
|  | H7/R2 | 1.486±0.216 | 2.2519±0.5384 |
| BBB transcytosis | F12/R2 | 1.960±0.180 | 3.658±1.104 |
|  | H7/R2 | 1.751±0.155 | 3.551±1.034 |

As a complementary experiment in the chip model aiming to confirm the transcytosis result, we perfused a single tube with equimolar amount of the targeting Nb (F12 or H7) and control Nb R2, which were labeled with different fluorophores, respectively **(Fig. 6e)**. This allowed each tube to serve as its own control. Importantly, this method minimizes the need to access tube leakage during the transcytosis assay by using the dextran tracers. As fluorophores with different spectrums cannot be directly compared based on fluorescence intensity, we introduced an index termed “**trans/assoc**”, which could show how a Nb is prone to transcytose the brain endothelial tube. This index represents Nb association per neural cell from bottom channel normalized to Nb association per endothelial cell from tube channel, hence, the index values can be directly compared between Nbs labeled with different fluorophores. As a dual-control Nb group where Alexa Fluor-647 R2 and Alexa Fluor-488 R2 were co-administrated in the tube lumen, their **trans/assoc** index values were found to be close. This proved that the Alexa Fluor-488 R2, when co-administrated with other Alexa Fluor-647 labeled Nbs, can serve as a proper control. As a result, we found that the **trans/assoc** values of F12 and H7 were both significantly higher than the co-administrated R2 **(Fig. 6e)**, indicating actual tube transcytosis of both the Nbs.

## Discussion

### ECM gel should reach an optimal stiffness to achieve a reliable NVU model

Endothelial tubes growing against 4 mg/ml collagen gel marks a standard protocol in the OrganoPlate [46, 47]. Notably, it is important that collagen concentration should reach a balance between gel longevity and in-gel cell compatibility. As shown here by us and others, collagen gel will be gradually degraded by the growing endothelial/epithelial cells (e.g., by matrix metalloproteinases [62]). Along with gel degradation, the Phaseguide structure of the OrganoPlate emerges as an abrupt obstacle to the expended tubes **(Fig. 4g)** [46, 47, 51, 62, 84, 85]. This led to a physiologically aberrant tube morphology which could potentially compromise tube properties. Moreover, the tube expanded in diameter as collagen decomposed, causing a decreased shear stress in the lumen thereby alleviating its beneficial effects on the endothelial cells. A possible approach to tackle this is to increase the collagen concentration which could slow down gel degradation. Even though collagen as high as 7-8 mg/ml provides superior stiffness for the endothelial tube and meanwhile still permits the survival of the embedded glia cells [42, 58, 86], the neurons were nonetheless only able to survive at a collagen concentration below 2.5 mg/ml [36, 42, 87–89]. Therefore, the neurons had to be seeded in the bottom channel of the chip using a softer neuron-friendly gel. As an optimal combination, we used 5 mg/ml collagen gel for the middle channel and 5 mg/ml Matrigel for the bottom channel. This double-gel solution yielded better tube stability and provided an optimal ECM environment for culturing the astrocytes and neurons. Importantly, the 5 mg/ml collagen allowed the neurons to directionally extend their neurites (mostly axons) into the gel while confining their somas in the Matrigel. We found the orientated neural outgrowth was independent of the tube presence as this phenomenon can also be seen in tube-free chips **(Fig. S8)**. Even though endothelial cells can secrete neurotropic factors [90], we found that the axonal outgrowth became less pronounced after tube seeding. Thus, the orientated neurite outgrowth seemed to be more a consequence of the medium perfusion in the top channel of the chip that efficiently created neurotrophic cytokine gradients across the gel [91, 92], whereas this process was partially blocked by the presence of the tube. Notably, the structure of axonal-vascular contact allows for studying the bi-directional transmission of misfolded protein aggregates in the NVU model. For example, in synucleinopathies such as Parkinson’s disease (PD), pathological α-synuclein aggregates contribute to disease exacerbation by not only spreading between neural cells [93–95], but also by transporting bi-directionally across the BBB [96, 97]. Furthermore, the segregated and directed axonal outgrowth, similarly to other reports [91], could provide an extra platform to model the neural projection pathway. As is known, retrograde axonal degeneration is one of the hallmarks of PD where the dying of nigral dopaminergic neurons starts from their distal terminals that innervate the striatum [98]. The modeling of such structures will help us better understand how these neurodegenerative diseases progress and guide the development of effective therapeutic interventions.

### Barrier integrity as an outcome of shear stress, gel stiffness, medium additives, and co-culture

An impermeable endothelial barrier is required for BBB transport studies. Here we will discuss different culture parameters that could affect tube integrity in the OrganoPlate model.

Serum is important in maintaining endothelial phenotypes including the barrier integrity [60]. In our early study, we have shown that proper serum level (e.g., 5%) is important in maintaining optimal barrier tightness of the brain endothelial layer that grows on the physical membrane (e.g., the transwell insert) **(manuscript under review)**. However, in the current model where the endothelial tubes grow directly against the ECM gel, serum of such level in the endothelial medium induced intense tube expansion which caused faster decomposition of the ECM gel and hence a greatly reduced model longevity. Importantly, we found that reducing the serum level to 2% could greatly prevent the tube expansion whereas this did not substantially compromise tube integrity.

Flow-induced shear stress is a major factor for the greatly promoted BBB integrity in the microfluidic models [99]. Shear stress upregulates the expression of various junctional proteins, which subsequently increases barrier trans-epithelial electrical resistance (TEER) [25, 26, 41, 44, 46, 48, 100, 101]. It was found that shear stress of the cerebral microcirculation was estimated to range from 0.01 to 10 dyn/cm^2^ in capillaries and 10–100 dyn/cm^2^ in arterioles [102]. However, shear stress created in the OrganoPlate system was reported to be about 1.2 dyn/cm^2^, which was much lower than that experienced by the blood vessels of similar diameter and curvature from *in vivo* [46, 47]. Some *in-vitro* studies showed that only 10-20 dyn/cm^2^ could substantially promote the integrity of hBMEC-derived tubes [26, 58]. Therefore, it is argued that the improved tube impermeability in the presence of medium perfusion in the OrganoPlate could have been the benefit of improved supply of oxygen and nutrients rather than shear stress [46, 103]. On the other hand, we have noticed that the medium flow induced by periodical rocking of the MIMETAS rocker remains flowing for only 2 min within the 8-min rocking cycle. Adjusting the rocking interval to 2 min/cycle was able to achieve a non-stopping flow but this led to decreased tube integrity over time. Such compromised barrier integrity might be explained in two ways. Firstly, the bi-directional flow has already been considered as an aberrant flow type which is detrimental to the endothelial cell morphology and barrier integrity [30, 104]. Thus, a more frequent reoccurring of the bi-directional flow was supposed to exert more negative effects to the tubes. Secondly, the increased periodical oscillation of the OrganoPlate could also spatially affect the endothelial cells’ behaviors. In addition, it remains unclear if this spatial unsettlement will also interfere with the behaviors of the neural cells, especially with the neurite outgrowth. Apart from shear stress, hydrostatic pressure (or transmural pressure) is another critical parameter that modulates barrier functions [21]. Hydrostatic pressure improves barrier integrity by driving the mobilization of intracellular TJ proteins more towards the cell borders without causing substantial increase in their expression [105]. Besides, hydrostatic pressure is important for maintaining stable morphology of the tubes and prevents endothelial sprouting into the surrounding gel [58]. In microfluidic models, higher hydrostatic pressure can be created by increasing the depth of the medium column [21, 58]. Collectively, shear stress and hydrostatic pressure play complementary roles in maintaining optimal vascular impermeability. It is conceivable that a syringe-based medium reservoir module [44] or pneumatic pump system [29, 106] may be developed for the OrganoPlate platform to completely overcome the problem of the plate oscillation, periodical flow, and low shear stress and hydrostatic pressure.

Barrier integrity is also profoundly influenced by the material type and stiffness where the endothelial cells grow on. Endothelial barriers growing on the surface of the ECM gels showed a much lower integrity than those growing on an artificial membrane such as the transwell membrane [59, 107]. For filter-free featured tubular models, increasing gel concentration producer higher gel stiffness [75, 86, 88, 89, 108–110], which in turn yields better tube stability and tube tightness [30, 74, 110]. In OrganoPlate-based models, we found that 7 mg/ml is the highest concentration for the collagen to be still injectable in the channels. Tubes growing against 7 mg/ml collagen gel showed better tube morphology and barrier integrity than those growing against 5 mg/ml gel **(data not shown)**. Nonetheless, it should be noticed that hydrogel exhibited reduced mechanical properties (e.g., modulus, ultimate strength, and peak strain) over prolonged incubation time [111]. On the other hand, denser gel leads to increased fiber density and smaller pore size after polymerization, which in turn hinders nutrient exchange and the diffusion of molecules to be tested in the gel [108]. It has been reported that the passive diffusion for full-size IgG was still permissive in collagen gel of 7-8 mg/ml [58]. However, the diffusion of larger therapeutical molecules remains to be checked under such high gel concentration. We also compared tube integrity between 5 mg/ml collagen gel with or without genipin crosslinking and found that the tubes growing against crosslinked gel displayed significantly better barrier tightness and morphology. It was also reported that genipin crosslinked collagen stabilizes microvessels under low shear stress or low hydrostatic pressure [112]. However, similar to using a denser gel in the model, crosslinked collagen was not suitable for modeling neural-vascular interactions or for modeling BBB extravasation of the immune cells as it totally blocks cell entry. Alternatively, other crosslinking methods can be used to bypass the drawback of the genipin approach, such as the glycation method [113].

Cytokines secreted by the neural cells and the direct neural-vascular contact are both important signaling mechanisms to regulate BBB functions and barrier integrity [1, 2]. In our NVU chip model, we have seen extensive neurite outgrowth and astrocytic migration in the gel which subsequently formed broad contact with the endothelial tubes. However, tube integrity was not improved in this case. Interestingly, several microfluidic NVU models demonstrated as well no further improvement in tube integrity by neural co-culture [29, 42, 46]. A possible explanation is that the astrocytes were not in a high number to produce enough soluble factors as well as to form enough endothelial contact that could substantially improve the barrier integrity [99]. We tried to seed more astrocytes in the model, only to find a faster speed for these cells to contract the collagen gel. Another reason could be too many neurons in the model. Even though the culture of high amount of LUHMES neurons produced extensive neural-vascular contacts without contracting the collagen, we noticed this led to a faster pH lowering of the neural medium, as was observed by frequent neural medium yellowing. Consequently, metabolic waste of these cells, including the resultant low environmental pH as well, could be a potential disruption for the endothelial barrier. Furthermore, cellularization of the ECM gel was shown to weaken its mechanical properties [109]. Therefore, optimization of the neural seeding density and actions to eliminate undifferentiated neuron progenitor could improve the model [30]. Importantly, we also realized that the increased model complexity, mainly contributed by the middle channel collagen gel that not only supports the tube without a physical membrane but also causes the random and uncontrollable in-gel outgrowth of the neural cells, has led to the decreased model reproducibility as well as batch-to-batch variations.

This was partially revealed by the different appearances of each co-culture chip **(Fig. S9)**, and also by the more scattered data points in the BIA **(Fig. 4f)**. Similarly, a decreased consistency in the barrier permeability was also demonstrated in a BBB model when the medium flow and glia co-culture were applied [29]. Together, it is suggested that reaching a balance between model reproducibility and model complexity is important for developing the NVU models.

### Modeling neuroinflammation in OrganoPlate NVU model

Disruption of the BBB can be either the cause or consequence of neuroinflammation, a hallmark in various neurological diseases [3, 77, 79, 80, 114]. Previous studies have extensively investigated the immune responses of brain endothelial cells by exposing inflammatory cytokines to the apical side of the endothelial layer, which represents the way to model BBB damage of an exogenous source such as systemic inflammation/infections [30, 40, 115–121]. On the other hand, for neurodegenerative diseases that have an endogenous etiopathogenesis such as genetic mutations in the neural cells, neuroinflammation influences the BBB primarily on its basal side [3, 40, 79, 122–126]. In the NVU models, this can be achieved by incorporating mutation-carrying neural cells [43] or disease inducers at the basal side of the brain endothelium [33, 40, 119]. Here, we showed that modeling neuroinflammation with endogenous causes is possible by administrating inflammatory cytokines at the bottom channel of chips. Immune responses of the endothelial cells, including decreased expression of junctional proteins, compromised barrier integrity and recruitment of circulating immune cells, were comparable between that when inflammatory cytokines were given to the tube lumen or to the bottom channel. Noteworthy, inflammation at the “brain side” was supposed to induce a more sever endothelial damage as the inflammatory cytokines not only would directly undermine the endothelial cells but could also inflame the glia cells (i.e., astrocytes and microglia) to synergistically release more inflammatory cues [40]. However, as we did not include microglia in the model, we were unable to reproduce the inflammatory communications between astrocytes and microglia, which triggered cascade reactions and further escalated the BBB damage [76].

Alignment is a common mechanobiological response of the endothelial cells [104]. However, whether medium flow aligns the *in-vitro* cultured hBMECs remains controversial [26, 29, 44, 127, 128]. One study showed that the shear stress must reach 10-20 dyn/cm^2^ to align the hBMECs [26]. Another study pointed out that low density of the endothelial cells is key to drive their alignment against the flow as high cell density might reduce the migration rate of the cells [29]. Coincidentally, in our study, hBMECs alignment was observed after TNFα and IL-1β exposure, the treatment of which induced cell death and consequently reduced cell density. This phenomenon has been also reported by others [81].

### Medium flow is important for drug screening

Medium flow in the microfluidic models has greatly revolutionized the field of *in-vitro* drug screening. Medium perfusion, which mimics the blood stream in the body, could greatly alleviate nonspecific association of the tested molecules to the endothelial cells [15]. Moreover, flow condition even enables the researchers to test how the shape of nanoparticle-based drugs influences their cell binding properties [129, 130]. In this study, we demonstrated the more specific targeting performance of a BBB-targeting Nb in the OrganoPlate-based model. Compared to the transwell model, medium flow in the OrganoPlate could significantly decrease the nonspecific interaction between the negative Nb control R2 and the hBMECs. In parallel, binding of the active-targeting Nbs F12 and H7 to the hBMECs decreased as well under flow condition. In line with our previous finding, this indicated that for Nbs that have specific targets on the cell surface, a portion of them could still be endocytosed/transcytosed in a nonspecific way when incubated statically on the endothelial cells **(manuscript under review)**. Quantitative comparison of transcytosed Nbs between the two models is barely possible as their totally different geometries and methods bring great discrepancy to data analysis and data comparison [107, 131]. By processing the data in a fold-change manner within each model, we were able to have them fairly compared. As a result, it was found that the Nb transcytosis was more pronounced in the microfluidic NVU model. Notably, even though the medium flow greatly reduced the nonspecific association of all tested Nbs, it amplifies the gap in the transcytosis efficacy between the real positive ones and the negative ones. Thus, by using a universal negative control across different model formats, data resulted by our data processing method indicates the NVU chip model is more faithful in revealing the true effectiveness of the candidate drugs in targeting the BBB.

### Limitations in the current OrganoPlate NVU model

Even though the three-channel architecture gives the OrganoPlate high flexibility in designing tubular models, the double-gel design of our NVU model could however be improved by a fourth channel [42, 43]. This additional channel is to be placed below the Matrigel channel in order to facilitate the medium nourishment and waste exchange of the neural cells embedded in the gel. More importantly, the fourth channel could make it possible to retrieve medium sample from the “brain side” during BBB transcytosis assay, which allows for calculating exact Papp value for the tested molecules and thus expanding the drug screening application beyond the fluorescently labeled ones. Nonetheless, as a more realistic solution, the choice of an optimal ECM gel capable of supporting both the tube integrity and the neural cell growth within it could make the NVU model fit the 3-lane chip perfectly. Indeed, a blended ECM gel consisting of 2 mg/ml collagen, 1 mg/ml hyaluronic acid and 1 mg/ml Matrigel was reported to not only allow for more extensive astrocyte processes outgrowth in the gel but also yield a much higher gel stiffness than that of 4 mg/ml collagen [86]. More importantly, reducing collagen concentration was found to keep the embedded astrocytes at a more quiescent state, which mimics the *in-vivo* situation [86]. Another hybrid ECM gel combining 3.7 mg/ml collagen and 7.1 mg/ml hyaluronic acid was documented [30]. Despite the high stiffness of this hybrid gel which guarantees the growth of a tight tube, it also grants the growth and differentiation of neurons inside of it [30]. Furthermore, one group set up the tri-culture of neurons/astrocytes/microglia in a commercial ECM gel named BME type R1 [44]. Due to enrichment of entactin which connects laminins and collagens, the BME type R1 gel yields a reinforced hydrogel structure which was able to firmly support the tube formation. These hybrid gels are worthy of exploring in the OrganoPlate system.

We are currently missing two important cell types in the model which are pericytes and microglia. Notably, pericytes were found to be more important in inducing a tight endothelial barrier as compared to astrocytes [99]. Besides, both pericytes and microglia are involved in neuroinflammation-associated BBB alterations [132, 133]. Given that there are complex crosstalk among different cell types within the NVU [1, 2, 76], it necessitates a complete-cell-type composition for faithfully modeling the NVU under both physiological/pathological conditions [30, 33, 40].

### Perspective on the use of hiPSCs in NVU modeling

Due to limited availability of NVU-related cell types of a human origin, there is an increased demand of using human induced pluripotent stem cells (hiPSCs) to derive these cells. Importantly, the use of patient iPSCs provides a chance for the development of disease-specific models [134, 135]. Therefore, the use of hiPSCs in combination with microfluidic technology, could contribute to a more comprehensive NVU modeling and advance our knowledge in inherited neurological disorders as well as novel drug discovery. As a flanking work of this study, we also generated a hiPSC-derived brain endothelial tube mode in the OrganoPlate **(Fig. S10)**. Compared to the hBMEC tubes, we found that the iPSC-tubes showed significantly better barrier tightness. In parallel, a number of iPSC-derived BBB tube models were reported to achieve a physiologically relevant barrier integrity [25, 40, 101, 136]. The presence of high barrier integrity and the expression of extensive BBB-specific markers has made the iPSC-derived brain endothelial cells an optimal cell source for BBB/NVU modeling [134, 137]. However, lacking standardized differentiation protocols for these iPSC-derived brain endothelial cells makes the resultant models inconsistent in the phenotypes and biological behaviors [48, 67, 134, 138, 139]. More importantly, with current protocols, most iPSC derived brain endothelial cell cultures harvest cell contamination of epithelial genotypes [138, 140]. Incorporation of these epithelial cells in the model could lead to mistaken data and conclusions. As possible solutions, it was found that the presence of 3D gel culture scaffold, shear stress and neural co-culture were able to decrease the epithelial identity and promote the iPSC-derived brain endothelial cells towards more authentic BBB phenotypes [25, 67, 139, 141]. These findings also support the hypothesis that microenvironmental cues play an important role in the region-specific phenotyping of the endothelial cells [138, 142–145].

## Conclusion

In this study, we optimized the protocol of the 3-lane OrganoPlate to generate a full-3D NVU model. Our model moves one step further in reproducing the complex microstructure of the NVU as compared to the transwell-based or other microfluidic-based models. It provides a platform for studying fundamental NVU functions *in vitro*, and for investigating endothelium disruption and immune cell extravasation upon neuroinflammation. Moreover, we showed the NVU model has better fidelity in screening for potential BBB shuttle molecules which could be further developed into promising brain-targeting drugs. Due to commercial availability, high throughput feature and compatibility with most used experimental devices, this OrganoPlate-based NVU model holds the potential to become a physiologically relevant BBB model with easy-to-use and high throughput drug screening features.

## Acknowledgements

PBMCs were a courtesy of the UMCU. We thank Suneel Narayanavari (NanoCell Therapeutics B.V.) for isolating them. We thank Alessia Di Maggio (Cell Biology, Neurobiology and Biophysics, Department of Biology, Science Faculty, Utrecht University) for constantly producing the Nbs.

## Competing interests

The authors declare that they have no competing interests.

## Supplementary figures

**Fig. S1.**
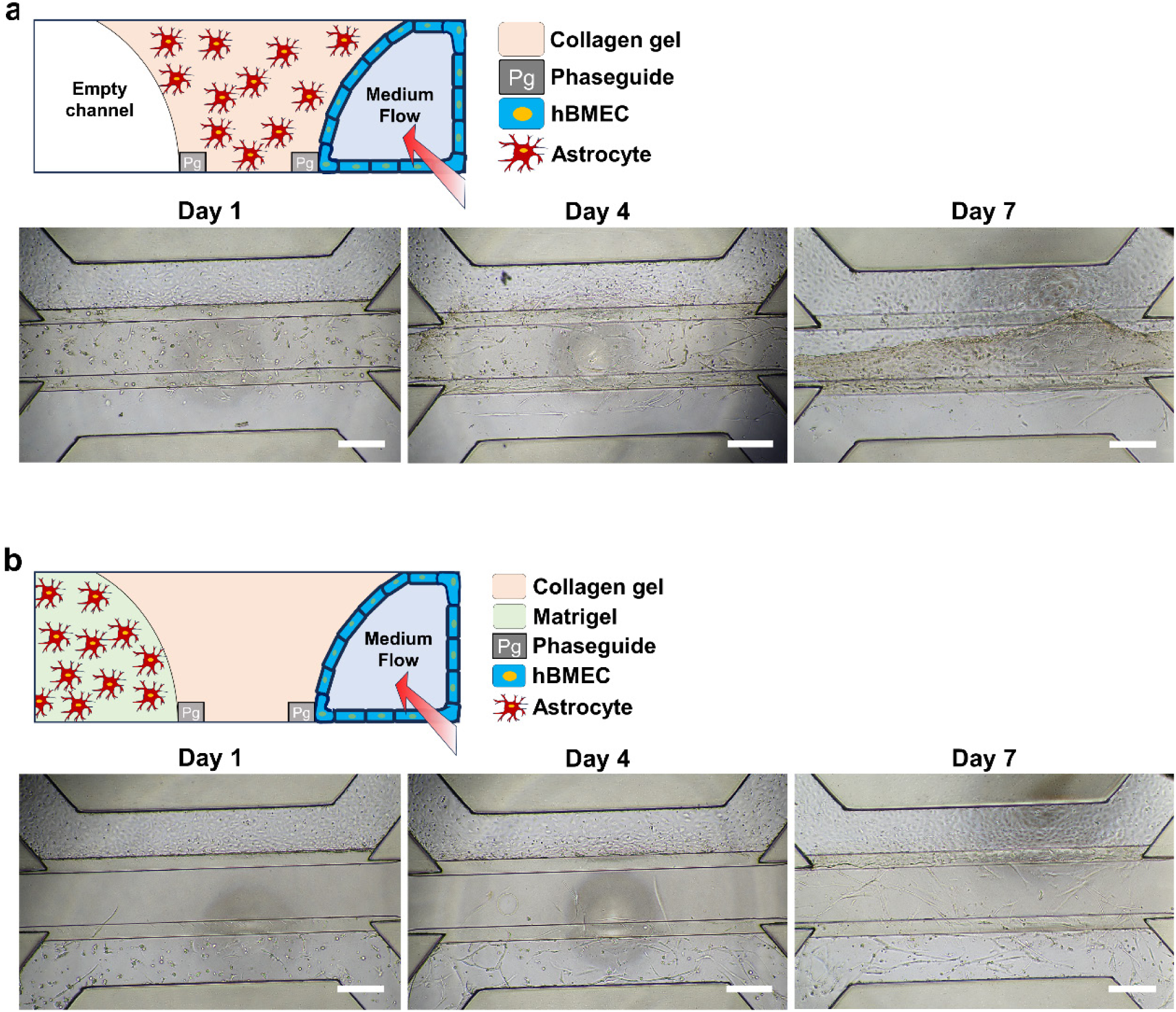
Screening for optimal astrocytes seeding strategies in the 3-lane OrganoPlate chips. **(a)** Astrocytes directly seeded in collagen gel in the middle channel of the chip. **(b)** Astrocytes seeded in Matrigel in the bottom channel of the chip. Images were taken as bright field. Scale bars are 500 µm.

**Fig. S2.**
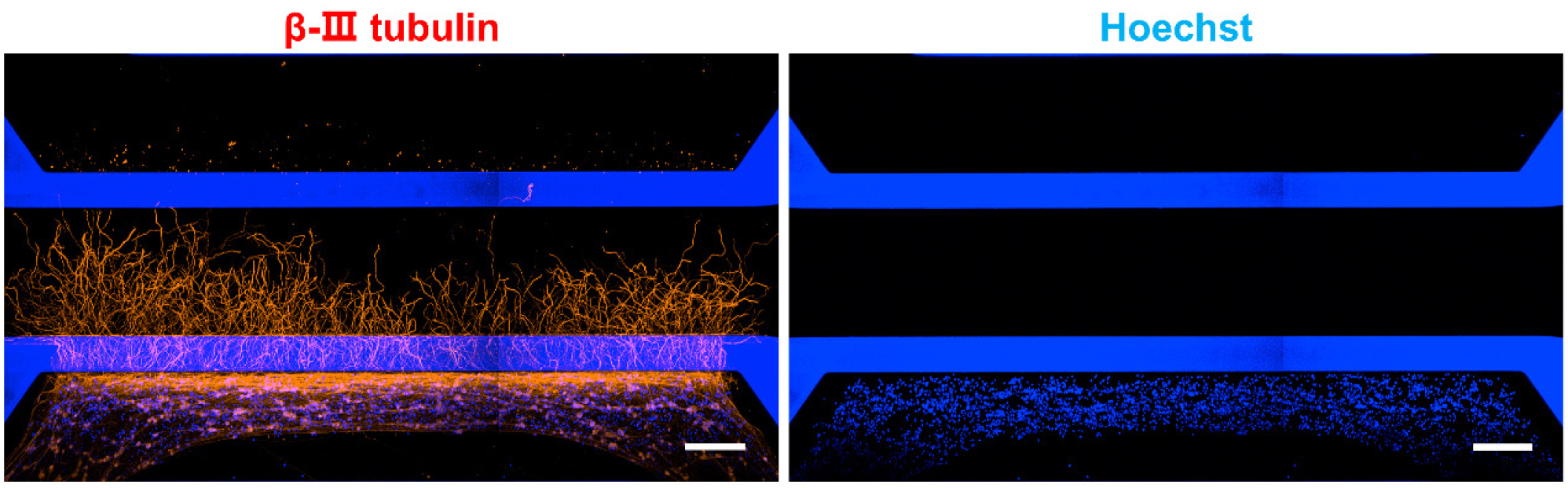
Neurites outgrowth of the LUHMES neurons in the 3-lane chip. Collagen gel was loaded in the middle channel. LUHMES neurons were seeded in Matrigel gel in the bottom channel. IFM images were taken 10 days after cell seeding. Scale bars are 200 µm.

**Fig. S3.**
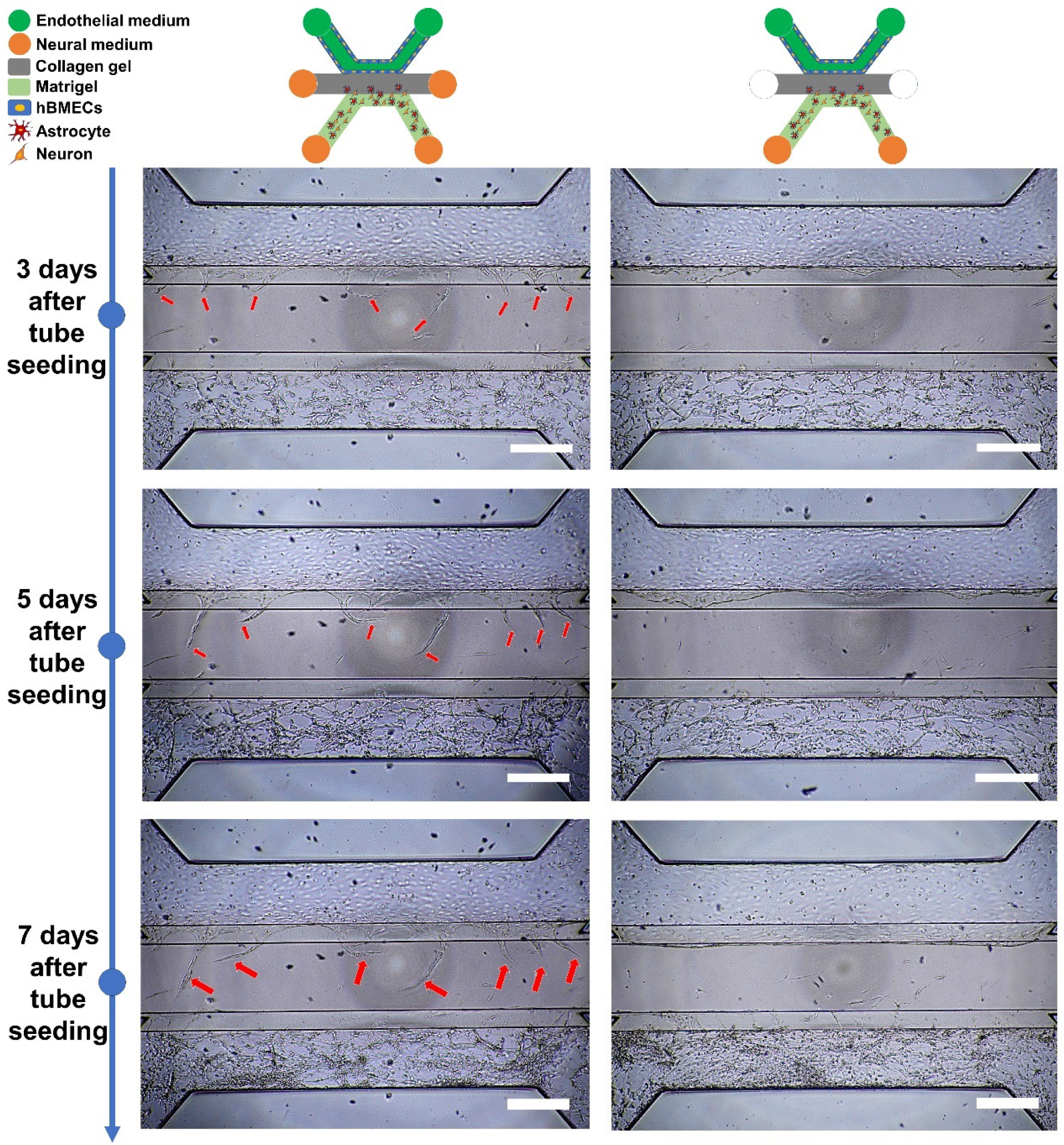
Neural medium given to the in/outlets of the middle channel of the 3-lane chips induces tube angiogenesis. Images were taken as bright field. Red arrows indicate the sprouts. Scale bars are 500 µm.

**Fig. S4.**
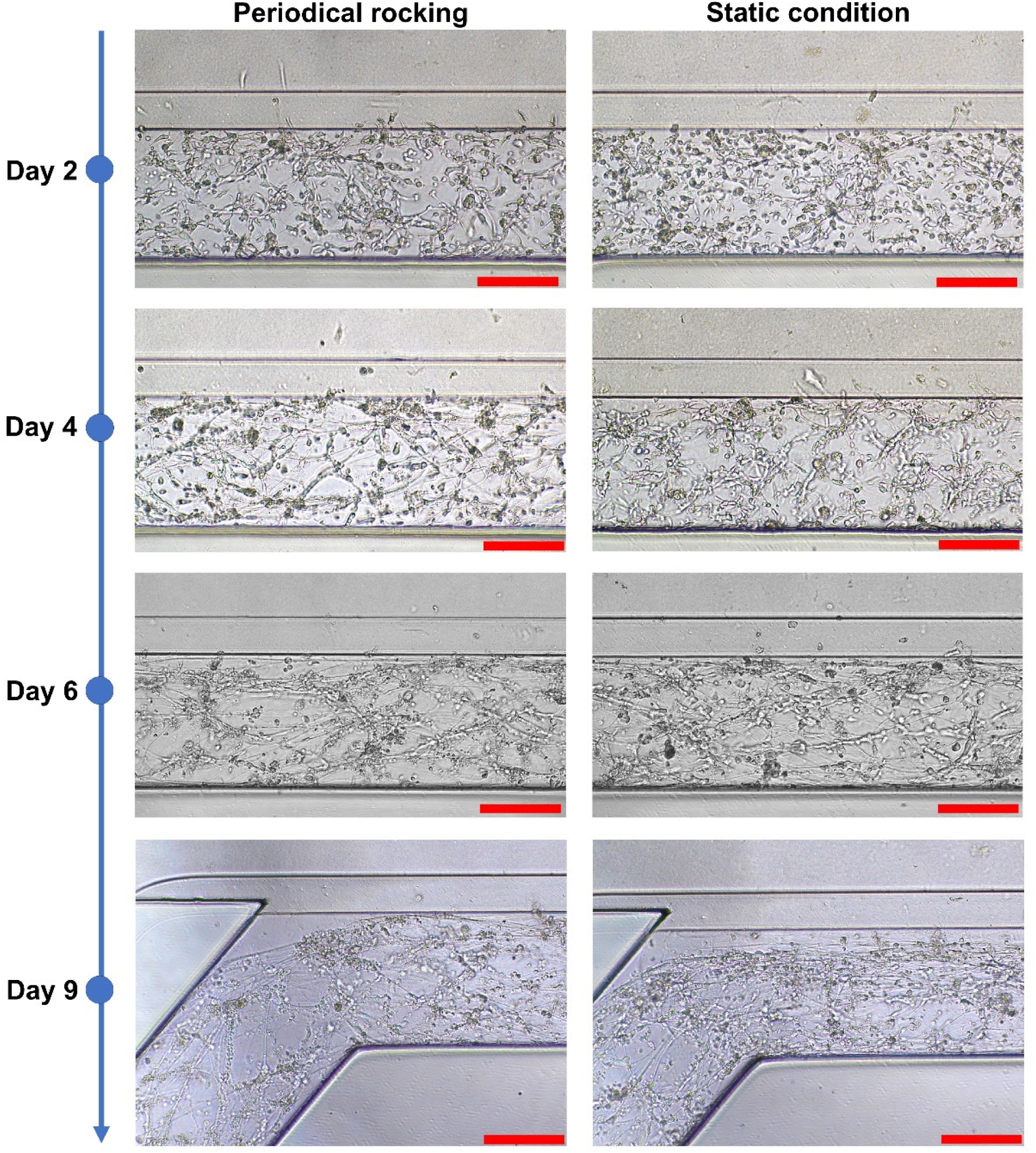
Impact of periodical rocking of the OrganoPlate on neural development of the LUHMES neurons. Images were taken as bright field. Scale bars are 200 µm.

**Fig. S5.**
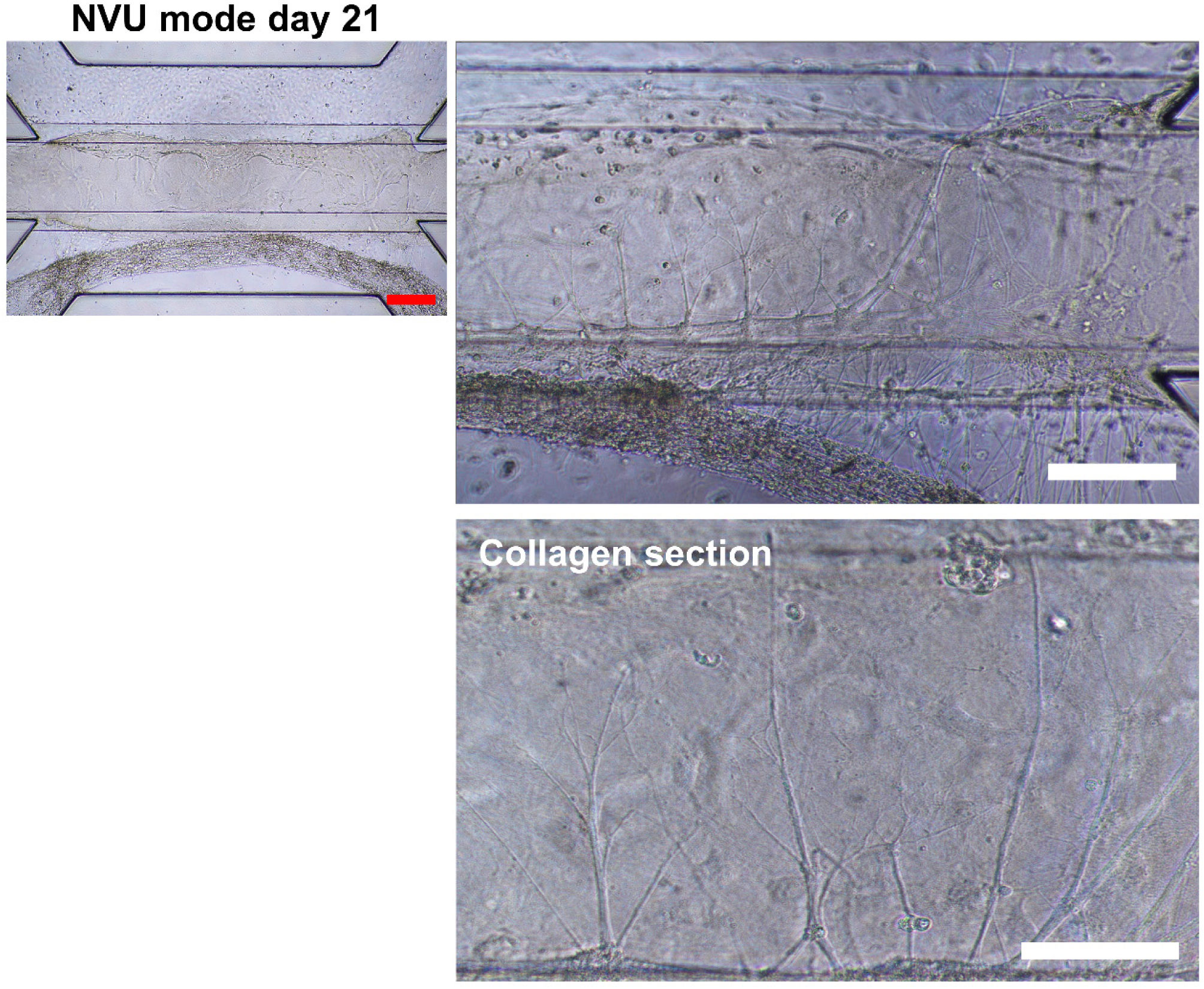
The NVU model after prolonged culture in the chip. Scale bars are 500 µm (red bar) or 100 µm (white bars).

**Fig. S6.**
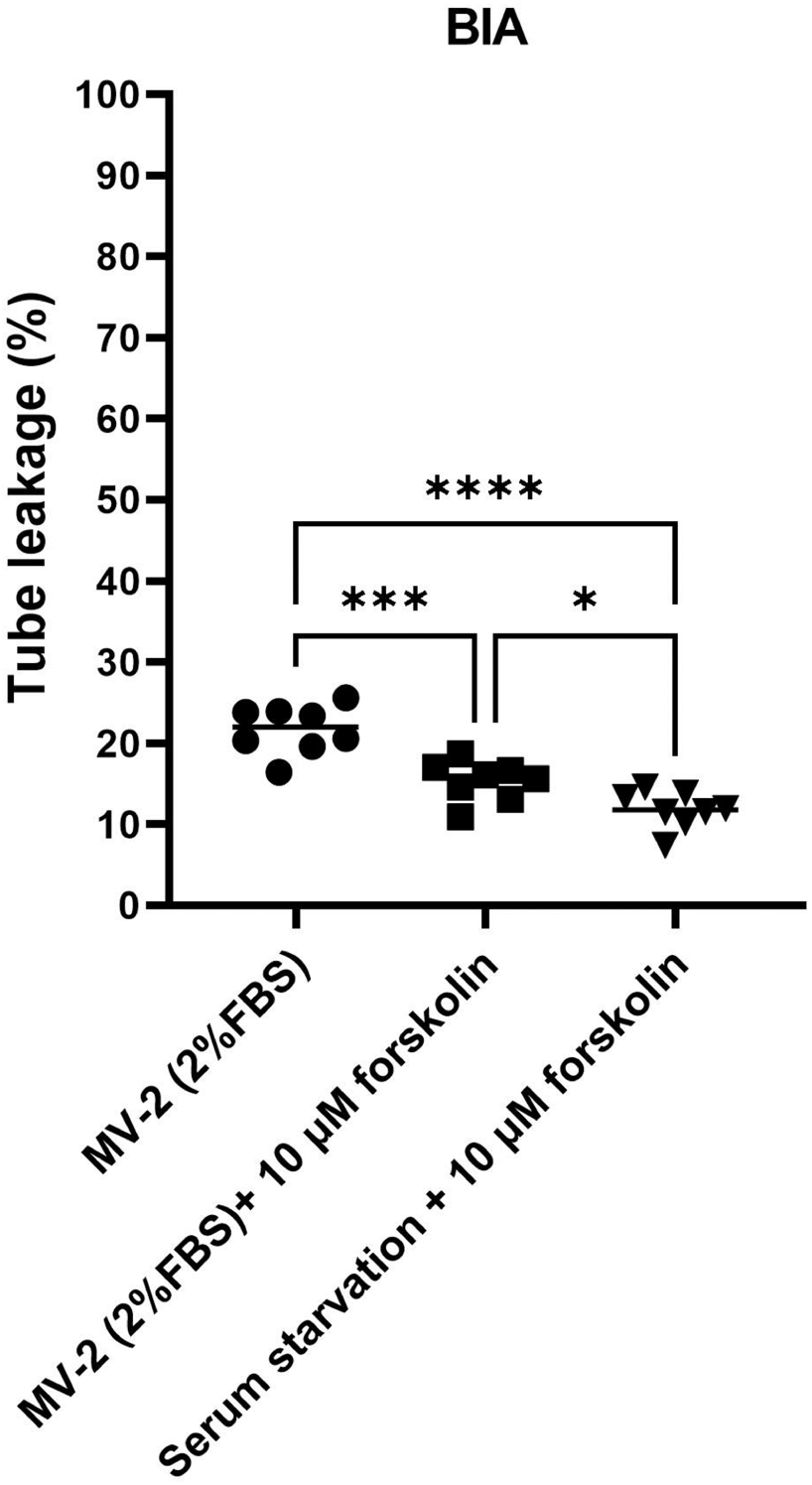
Effects of serum starvation and forskolin on the barrier integrity of the hBMEC-tubes. For quantification, 8 chips were used in each group. Graph shows mean ± SD. Statistical analysis was performed using one-way ANOVA followed by Tukey’s multiple comparisons tests; *P < 0.05, **P < 0.01, ***P < 0.001, ****P < 0.0001.

**Fig. S7.**
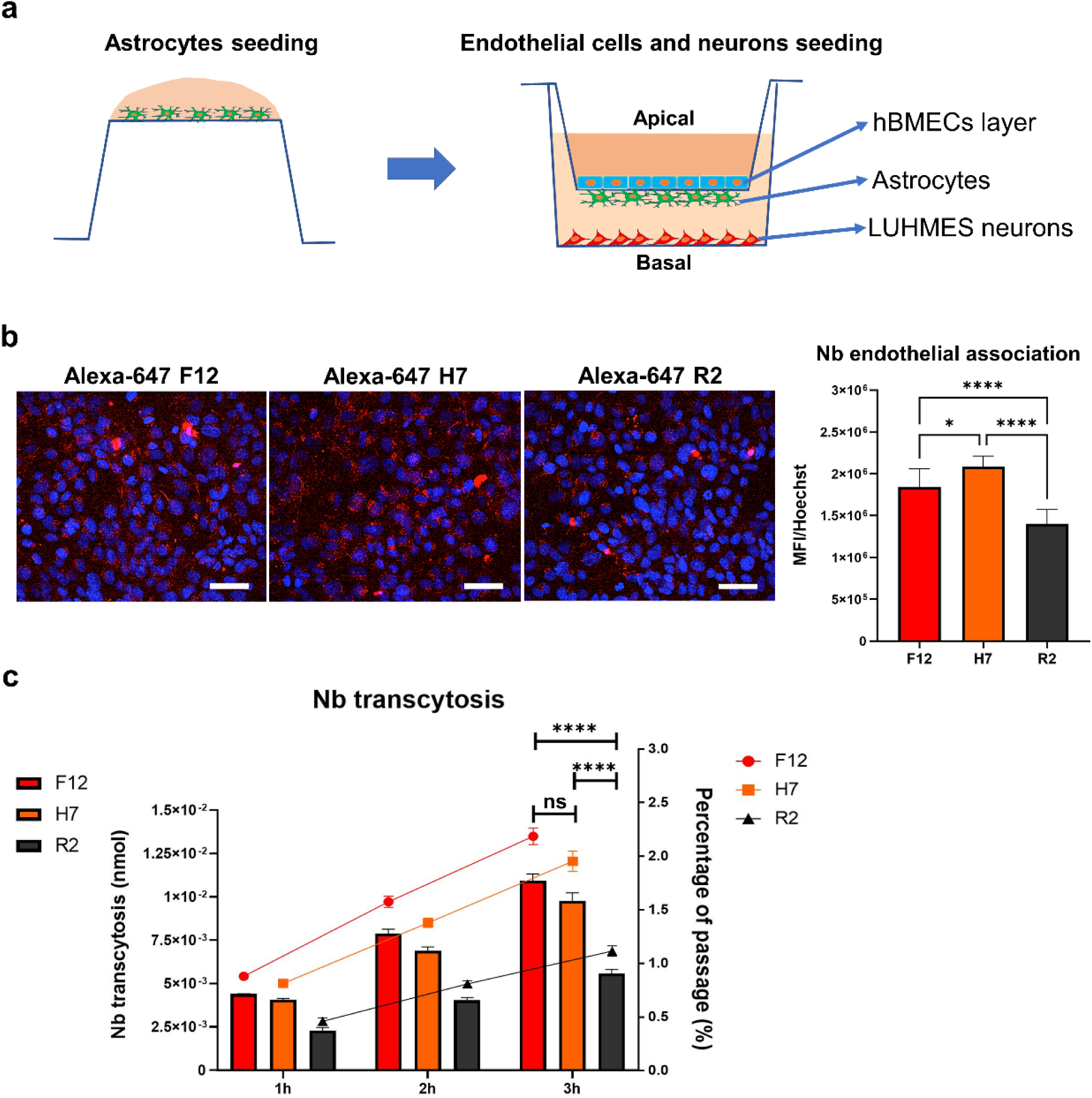
Nb transcytosis in the transwell-based NVU model. **(a)** Schematic protocol of the transwell NVU model. hBMECs grow in the insert as a monolayer. Primary astrocytes grow on the back of the insert. LUHMES neurons grow in the bottom of the outer well. **(b)** 1000 nM Alexa Fluor-647 labeled Nbs, where F12 and H7 target proHB-EGF and R2 targets Reactive Red 6, were incubated at the apical side of the endothelium. Endothelial association (binding + uptake) of the Nbs was evaluated by IFM after 3 h. A total of 30 fields taken from 3 transwell inserts were acquired for quantification. Fluorescence signal of the Nbs was normalized to counted cell number. IFM images were taken at maximum projection mode. Scale bars are 50 µm. **(c)** Alexa Fluor-647 labeled Nbs at 1000 nM were incubated at the apical side of the model. Medium samples were taken from the basal side and Nb transcytosis was evaluated by SDS-PAGE followed by fluorescence illumination. For quantification, each group contains 3 transwell units. All graphs show mean ± SD. Statistical analysis was performed using one-way ANOVA followed by Tukey’s multiple comparisons tests; ns indicates no significant differences; *P < 0.05, **P < 0.01, ***P < 0.001, ****P < 0.0001.

**Fig. S8.**
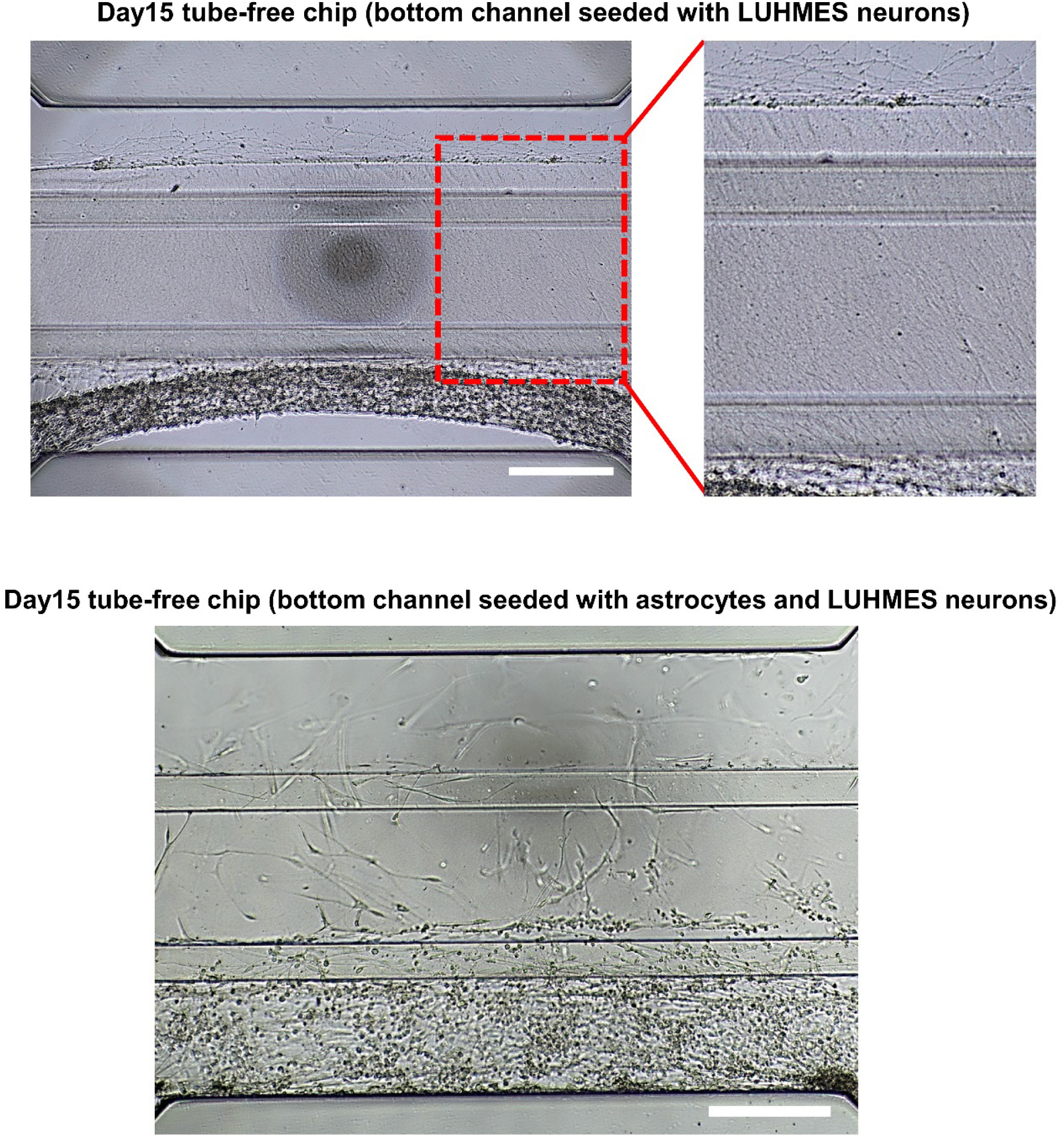
Oriented axonal outgrowth and astrocyte migration are independent of the presence of endothelial tube. Images were taken as bright field. Scale bars are 500 µm.

**Fig. S9.**
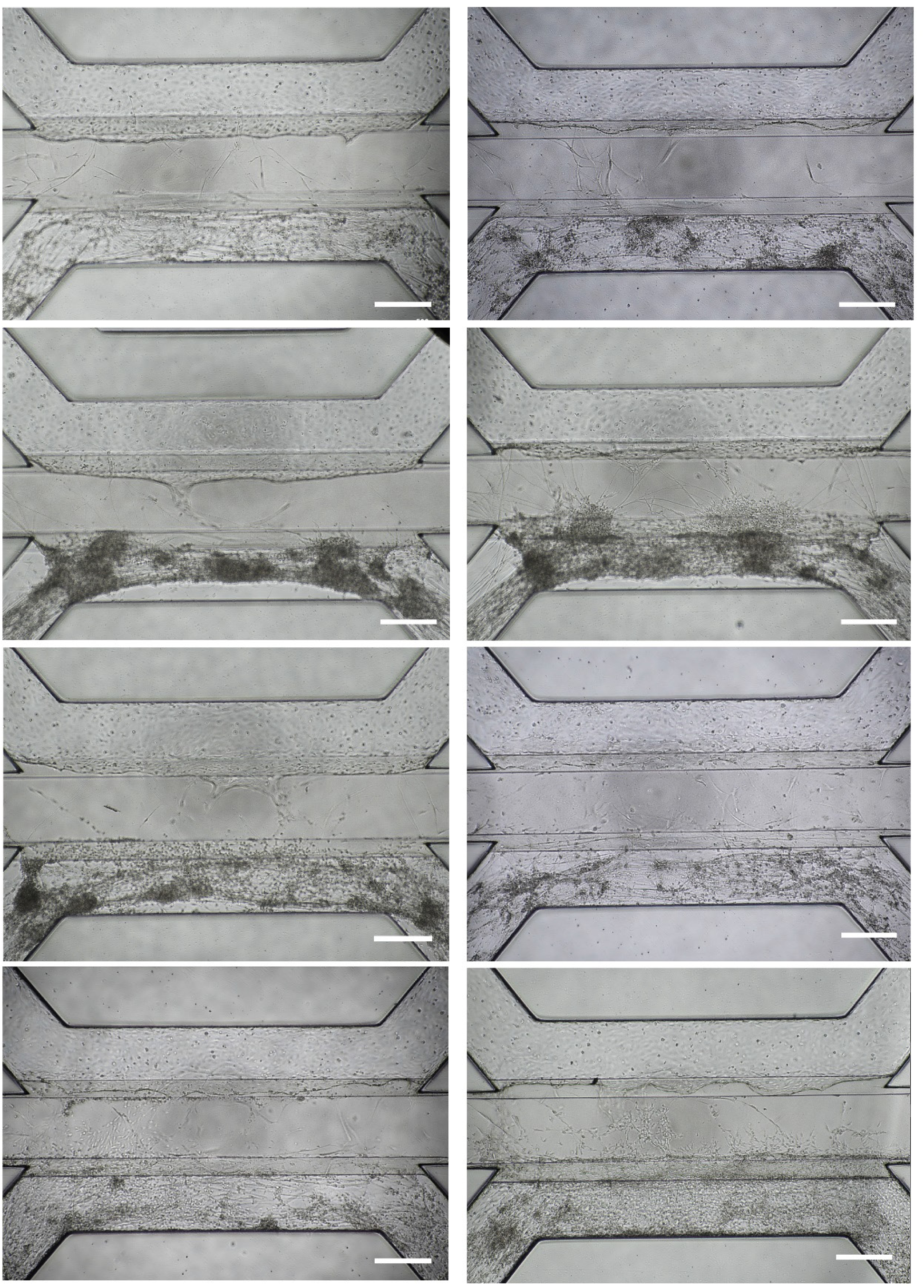
An example of quality inconsistency in tri-culture chips. Images were taken as bright field. Scale bars are 500 µm.

**Fig. S10.**
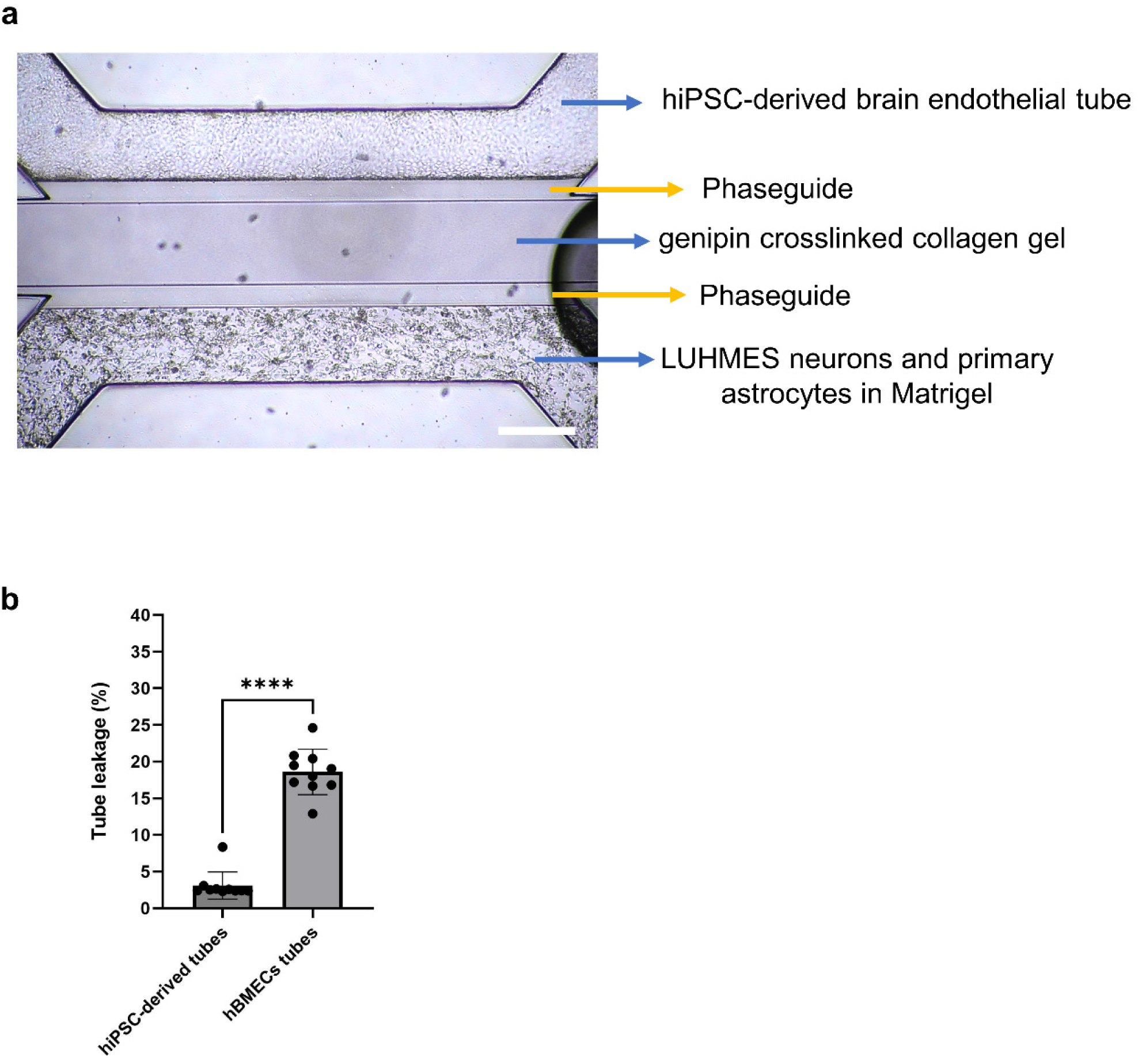
OrganoPlate-based NVU model comprised of hiPSC-derived brain endothelial tube. **(a)** Brightfield image of the model. Scale bar is 500 µm. **(b)** BIA between hiPSC-derived brain endothelial tubes and hBMEC-tubes. Each group contains 10 tubes. Graph shows mean ± SD. Statistical analysis was performed using student’s t-test; *P < 0.05, **P < 0.01, ***P < 0.001, ****P < 0.0001.

## Notes

### Competing Interest Statement

The authors have declared no competing interest.

